# Cell size-dependent mRNA transcription drives proteome remodeling

**DOI:** 10.1101/2025.10.30.685141

**Authors:** Dong Shin You, Christopher H. Bohrer, Purva H. Rumde, Ioannis Sanidas, Matthew P. Swaffer, Daniel R. Larson, Josh E. Elias, Michael C. Lanz, Jan M. Skotheim

## Abstract

Increasing cell size drives proteomic changes that impact cell physiology. However, the molecular basis of size-dependent proteome remodeling has remained unclear. Here, we develop an inducible Cyclin D1 expression system in human cells to generate populations of proliferating cells spanning over a two-fold size range. We use this genetic system to make comprehensive genome-wide measurements of mRNA and protein concentrations and stability. We find that protein and mRNA turnover rates are weakly related to cell size, but that mRNA concentrations are strongly size-dependent. This establishes that transcriptional regulation is the basis of proteome remodeling. Live-cell imaging of endogenous mRNAs using MS2 fluorescent protein binding motifs is used to measure how transcriptional dynamics change with cell size. Larger cells prolong transcriptional bursts and shorten inactive periods between bursts but maintain similar burst amplitudes to achieve transcriptional scaling. Taken together, our results show how transcription is modulated by cell size to remodel the proteome and alter cell physiology.

## Introduction

Cell size is a fundamental cellular characteristic that influences physiology, fitness, and fate. For example, the small size of red blood cells facilitates their efficient deformation and circulation,^1–3^ while adipocyte functions are modulated by changes in cell size.^4^ The importance of maintaining an appropriate size is seen in the striking uniformity of cell size within a given type as well as the strong association between aberrant cell size and pathological states including cancer, senescence, and aging.^5–15^ Yet, apart from avoiding more extreme ends of the cell size distributions, it is unclear why cells target a characteristic size.^16,17^

New understanding of the physiological importance of cell size came from studies examining the molecular mechanisms that link cell growth to division in order to control cell size. These mechanisms typically couple cell growth to the timing of G1/S transition and involve key cell cycle regulatory proteins. For example, as cells grow larger, they dilute inhibitors of cell division, like the retinoblastoma protein RB in humans,^18,19^ Whi5 in budding yeast,^20,21^ KRP4 in plants,^22^ and TNY1 in algae.^23^ Increasing cell size can also increase the concentration of cell cycle activators, as is the case for Cdc25 in fission yeast.^24^ The commonality between the mechanisms identified thus far is that they all involve size-dependent changes in protein concentrations that trigger cell division.^25^ The discovery of these size control mechanisms highlighted the fact that increases in cell size could drive changes to protein concentrations.

The discovery of size-dependent changes in the concentrations of cell cycle regulator proteins suggested that the concentrations of other proteins could also depend on cell size. This observation went against the prevailing paradigm that the concentration of the vast majority of proteins would reflect the total protein concentration, which is largely independent of cell size until cells become extremely large.^26–29^ However, recent work using modern proteomic technology has definitively overturned this paradigm. When proliferating mammalian cells are stratified by their natural size variation using FACS and analyzed with proteomics, the concentrations of individual proteins do not generally remain constant. Instead, proteome composition is globally remodeled as cell size increases.^30^ Such wide-spread size-dependent changes in proteome composition were found in all cells studied so far, including both eukaryotes and prokaryotes.^31–35^

Collectively, this work demonstrated that cell size must be tightly controlled because the further a cell deviates from its target size, the more compositionally different it becomes. The size-dependence of cell composition may explain the allometric relationship between cell size and biosynthesis. As cell size increases past its optimum, the concentrations of ribosomes and other biosynthetic proteins decrease, which coincides with a decline in growth rate and mitochondrial function.^36,37^ Ultimately, excessively large cells are unable to scale protein concentration with volume, leading to cytoplasmic dilution and senescence.^37^

A series of recent studies have shed additional insight on how excessive cell enlargement leads to senescence.^38^ The blockade of CDK4/6 activity, which drives cell enlargement by preventing cell division but not preventing cell growth, leads to p21 upregulation and permanent G1 arrest.^39^ The enlarged cells that do re-enter the cell cycle upon removal of CDK4/6 inhibition were more susceptible to DNA damage,^40^ potentially from the under-production of DNA repair machinery.^30,41^ These deleterious effects on genome integrity and proliferation were mitigated if cell growth during arrest was downregulated through the inhibition of mTOR or the PI3K pathway.^39–42^ Similar phenomena were found upon CDK7 inhibition and subsequent cell size increase.^42^ Importantly, the size-dependent proteome remodeling measured in normal-sized cells and the ultimate failure of excessively-large cells to scale protein biosynthesis with size are both consequences of a declining DNA-to-cell volume ratio, since neither phenomena occur when increase in cell size coincides with proportional increase in ploidy.^30,31,33,43^ These studies strongly support the conclusion that excessive cell growth during cell cycle arrest is a major contributor to cellular senescence.^14^

While increases in cell size drive important proteomic changes, the mechanism through which this happens is currently unclear. In principle, the size-dependence of any aspect of mRNA and protein synthesis and degradation could be responsible for the size-dependent changes to proteome composition. Here, we used a suite of high-throughput technologies to systematically measure major steps of the gene expression pathway to determine which, if any, are size-dependent. To do this, we first developed a genetic system that modulates the expression of *CCND1* in *RPE-1* cells to generate populations of cells that proliferate at different cell sizes. We find that human cells alter their transcriptome with the increase in cell size and that this change in transcriptome is likely driven by the mRNA synthesis and its altered bursting kinetics. By employing pulse SILAC-TMT and SLAMseq, we also find that both mRNA and protein turnover rates remain mostly stable and do not display differential changes across roughly twofold cell size. Our study pinpoints transcription as the key size-dependent pathway responsible for proteomic remodeling and provides insight into the potential molecular mechanisms involved in altered transcription dynamics.

## Results

### A new genetic system to probe how cell size influences mammalian cell physiology

To better understand how cell size influences mammalian cell physiology, we previously sorted proliferating cell lines by size in G1 using FACS and measured their proteome composition.^30^ Proteins whose relative concentrations increase or decrease with increasing cell size were termed super- and sub-scaling, respectively (**Figure 1A,1B**). In principle, the molecular mechanisms underpinning these size-dependent protein concentration changes could operate at the levels of transcription, translation, and turnover (**Figure 1C**). However, the necessity of sorting large numbers of cells limited our ability to rigorously probe each possible point of regulation using modern multiomics methods. To overcome this limitation, we sought to engineer a tractable experimental system to investigate how cell size impacts cell physiology.

**Figure 1.**
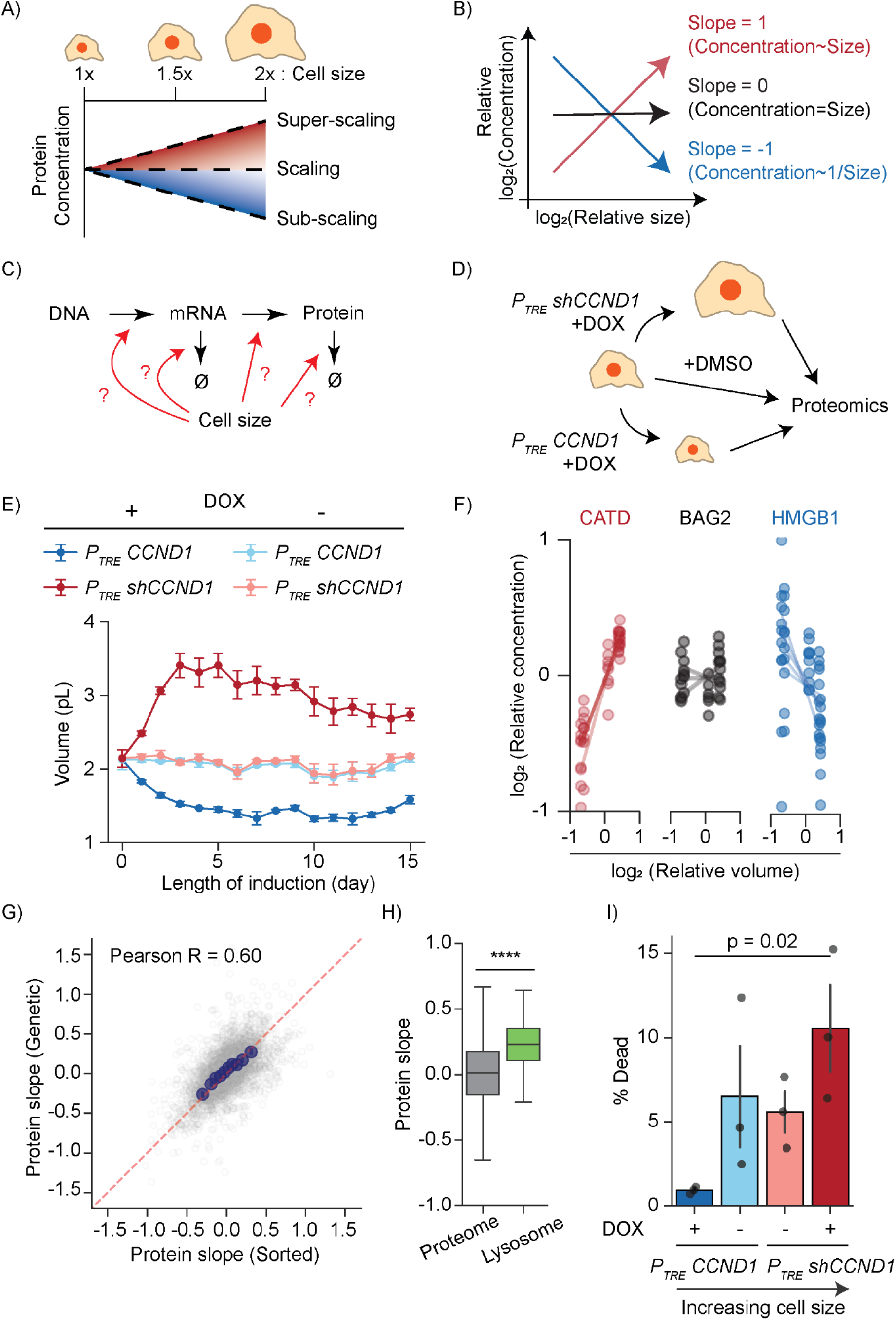
Size-modulation by *CCND1* manipulation remodels the proteome and impacts lysosomal vulnerability. **A)** Schematic summary of the scaling relationships between protein concentration and cell size as described previously in FACS-isolated G1 cells.^30,33,35^ **B)** A slope value quantifies how the concentration of a protein or transcript changes with cell size as previously described.^30,33,35^ Proteins with a slope of 0 maintain a constant relative concentration regardless of cell volume. A slope value of 1 corresponds to an increase in relative concentration that is proportional to the increase in volume, and a slope of −1 corresponds to dilution (1/volume). **C)** Schematic illustrating different aspects of synthesis and degradation that could impact the size-dependence of protein concentrations. **D)** Experimental strategy for obtaining differently-sized cycling cells using a doxycycline-inducible knockdown or overexpression of *CCND1.* **E)** Cell size dynamics for the indicated cell line after doxycycline induction. A new target size is reached by day 4 and mostly maintained for 10 days. Median cell volumes were measured using a Coulter counter and averaged from 3 separate experiments. Standard error of means are noted as error bars. **F)** Differential scaling of individual proteins. Relative protein concentration is plotted against relative cell size for three proteins that exemplify different size scaling behaviors - superscaling, scaling, and subscaling. Each dot represents an independent peptide measurement for the indicated protein. A linear regression is drawn from all the peptides detected for a given protein to find the slope value. A protein’s slope value is found by averaging all peptides’ slope values. For clarity, lower quality peptides were omitted from visualization. **G)** Proteome remodeling in genetically manipulated cycling cells (“Genetic”, this study) strongly correlates with our previous measurements in size-sorted G1 cells (“Sorted”).^30^ Each gray point marks the slope value of an individual protein and blue dots are the binned averages. At least 2 unique peptide measurements were required for inclusion in the plot (N = 3843 proteins). **H)** Proteins belonging exclusively to lysosomes (as defined in our previously published study^30^) superscale as a group relative to the entire proteome with cell size (p < 10^-5^). **I)** Larger cells are more vulnerable to lysosomal insults than smaller cells. After 5 days of size induction, cells were acutely treated with 3mM LLoMe then probed for cell death using Annexin V and cell permeability dye using flow cytometry. P-value shows outcome of unpaired t-test. Proportions of dying cells were found and the baseline death rate from control conditions were used to calculate death from LLoMe. N = 3 replicates.

To engineer a genetic system to study size-dependent expression, we took inspiration from the fact that deletion or overexpression of G1 cyclins can alter cell size.^44,45^ We therefore integrated a doxycycline-inducible system to titrate the concentration of a G1 cyclin, *CCND1*, in an hTERT-immortalized retinal pigment epithelial cell line (*hTERT RPE-1*, hereafter *RPE-1*). We chose to use these *RPE-1* cells as the basis of our system because they are diploid, genetically stable cells, and commonly used for cell size and cell cycle studies. To make larger cells, we added doxycycline to the media to express an shRNA that efficiently depletes cyclin D1. *CCND1* knockdown lengthens G1 phase and thereby increases cell size (*P_TRE_ shCCND1*). To make smaller cells, we also induce shRNA expression to deplete the endogenous *CCND1*, but now combine it with the overexpression of an shRNA resistant cyclin D1 allele (*P_TRE_ CCND1*) (**Figure S1A**) (see methods). This perturbation shortens G1 to decrease cell size (**Figure 1D,1E**). After 4 days of induction, the cells reach a new semi-steady state that persists for at least ten days, as measured by Coulter counter. Although the large cells show a gradual decline in size, possibly due to selection pressure for cells that can overcome the effect of *CCND1* knockdown, this decline is slow enough that it should not introduce major confounding effects into our analysis (**Figure 1E**). With the establishment of this tractable genetic system, we could now grow populations of cells that are cycling in steady state and have small, medium, or large sizes.

After engineering cell lines that cycle at different sizes, we sought to confirm that protein expression levels and their size-dependence were similar to those of unmodified cells that were sorted by varying sizes. To check this, we collected *P_TRE_ shCCND1* and *P_TRE_ CCND1* cells induced for 3,4,5, and 6 days. Uninduced cells were also collected as they had intermediate sizes. In total, we obtained proliferating populations of cells spanning just over a two-fold size range, and measured their proteomes using TMT mass-spectrometry (**Figure 1D, 1E**). A principal component analysis of our mass-spectrometry dataset showed that the proteomes of large and small cells also reach a relative steady-state after 4 days of induction, mirroring the time of stabilization of cell size (**Figure S1B, Figure 1E**). Next, for each individual protein, we determined its concentration’s size-dependence by calculating a protein slope value as described previously (**Figure 1F, Supplementary Table 1**).^30,33^ Briefly, the protein slope is derived from a linear regression of the logarithm of relative changes in cell size against the logarithm of the relative changes in protein’s relative concentration. A slope value of 0 means no change in concentration with an increase in cell size, while slopes of 1 and -1 reflect a two-fold increase and decrease in concentration with a two-fold cell size increase, respectively (**Figure 1B**). The slope values derived from our *CCND1* manipulation lines were highly correlated with those reported in our previous study using size-sorted immortalized and primary G1 cells (**Figure 1G, S1C, Supplementary Table 1**).^30^ The proteome slopes were also highly correlated with size-sorted primary mouse liver cells from another previous study (**Figure S1D**).^33^ Taken together, our analysis shows that *CCND1* manipulation provides a convenient and powerful platform to generate populations of cells cycling at different sizes, which can be used to study size-dependent proteome remodeling.

### Large cells are more vulnerable to lysosomal damage than smaller cells

While the size-dependence of some cell cycle regulatory proteins is functional, the vast majority of size-dependent proteome changes have not yet been linked to any function. In our original study of differential proteome scaling,^30^ we found lysosomal proteins to be among the most superscaling proteins (**Figure 1H**). Lysosomes are major signaling hubs for metabolism, and their biogenesis and function are critical in cellular proteostasis and aging.^46–48^ Because defects in lysosome function and increased lysosomal content are known markers of senescence,^49–51^ we wondered whether the super-scaling of lysosomal proteins reflects a size-dependent change in the relative importance of lysosome function. We hypothesized that if cells depend more on proper lysosome function with increased cell size, we would see increased death in large cells relative to small cells when we perturbed lysosome function. To test this, we inhibited lysosome function in our *CCND1* manipulated cell lines. After doxycycline-induction was used to reach the new target cell sizes (**Figure S1E**), cells were challenged with LLoMe, a lysosomotropic small molecule that physically compromises the lysosomal membrane and leads to cell death at higher concentrations. Upon acute treatment of 3mM LLoMe for 3 hours, we observed that large cells had higher rates of cell death than small cells as measured by Annexin V and cell permeability dye with flow cytometry (**Figure 1I**). Longer treatment for 48 hours with 50µM chloroquine, a small molecule that perturbs lysosomal pH and enzyme function, found a similar size-dependent increase in death, although the effect size was more subtle and relevant only in a narrow dosage range (**Figure S1F, S1G**). These size-dependent deaths were not due to a general vulnerability of large cells because treating cells with 20nM of the proteasome inhibitor bortezomib over 2 days resulted in smaller cells dying at lower concentrations, which was the opposite effect (**Figure S1H**). We therefore conclude that large cells are specifically more vulnerable to lysosomal damage, which may be due to the super-scaling nature of the lysosomal proteins.

### Cell cycle phase does not influence size-dependent proteome remodeling

Cell size and cell cycle phase are often confounding variables since cells that have spent more time progressing through the cell cycle are typically larger. In our previous work, we controlled for the potential confounding effects of the cell cycle by sorting differently-sized cells that were exclusively in G1 phase.^30^ Since most *RPE-1*s are in the G1 phase (∼70%), this facilitated the size-dependent sorting of G1 cells (**Figure 2A**, non-induced conditions). Size-sorting G2 cells is considerably more difficult since G2 cells account for only 20% of the population and less than 5% in the large population. In contrast, using our new genetic system we can directly sort all G2 cells after they have reached their new target sizes, making the process of measuring differently-sized G2 cells feasible.

**Figure 2.**
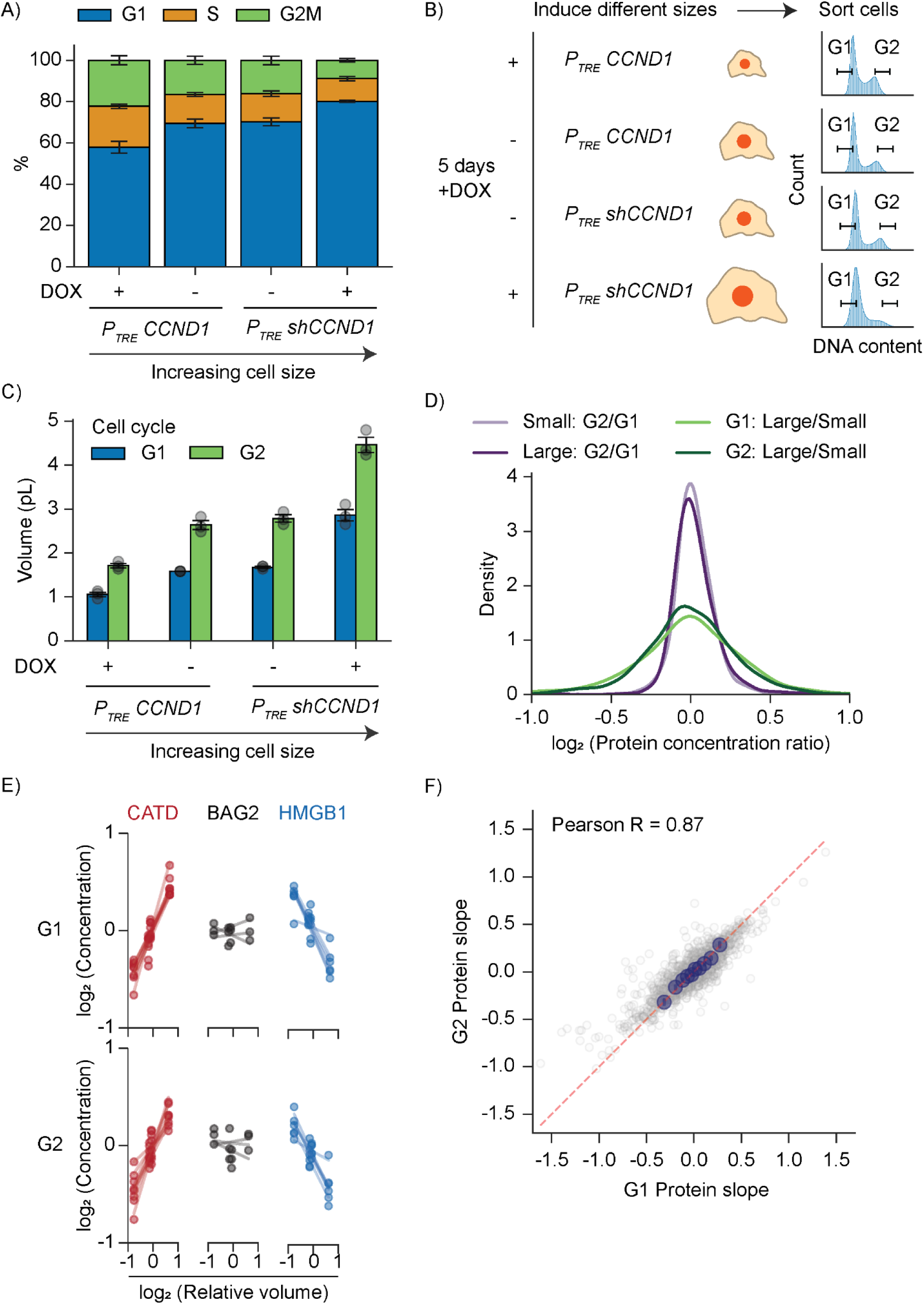
Cell cycle phase does not influence size-dependent proteome remodeling. **A)** Proportion of cells in different cell cycle phases after 5 days with and without doxycycline induction. Cells were stained with Hoechst and analyzed using flow cytometry for DNA content. Error bars mark the standard error of means from 3 separate measurements. **B)** Experimental design of cell cycle-dependent sorting of small, medium, and large cell populations. Cells were induced with doxycycline or DMSO for 5 days before staining by Hoechst. Cells in each of the indicated conditions were sorted by DNA content for G1 and G2 phases and analyzed by proteomics. **C)** Cell volumes of indicated cell lines after being sorted by cell cycle phase. Mean of the median cell volumes was calculated from 3 separate sorting experiments, with the points showing the raw median volumes. Error bars represent standard error of mean. **D)** Proteome changes caused by differences in cell size are larger than those caused by differences in cell cycle phase. Histograms are derived from the log_2_ protein concentration ratios between the indicated conditions. For example, “Small: G2/G1” is the ratio of protein concentration changes between G2 and G1 cells from the *P_TRE_* CCND1 +Dox condition. **E)** Differential scaling of the indicated individual proteins in size-sorted G1 and G2 cells (as in Figure 1F, each dot represents an independent peptide measurement for the indicated protein). **F)** Protein size slopes derived from size-sorted G1 and G2 cells. Blue dots are the binned averages. N = 1926 proteins.

To disentangle the effects of cell size and cell cycle phase, we generated cells of different sizes and then sorted by G1 and G2 cell cycle phase (**Figure 2B**). We confirmed that we were able to isolate cell populations based on cell size and cell cycle stage (**Figure 2C**) and then conducted TMT-mass spectrometry to probe their protein compositions. We find that the protein concentrations varied far more across cell size than cell cycle phase, which supports our hypothesis that proteome composition changes in large cells are mainly driven by size changes rather than changes to their cell cycle phase distribution (**Figure 2D**). Moreover, changes to the proteome were linearly correlated with cell size for both G1 and G2 sorted cells, meaning that both populations remodel their proteome consistently along the size axis (**Figure S2A**). After calculating the rate of change of protein concentrations with cell size and cell cycle phase as before (**Figure 2E, Supplementary Table 2**), we compared the linear slope of protein concentration changes between G1 and G2. We found that protein slopes derived from G2 cells strongly correlated with those from both G1 cells and those from our asynchronous populations of *CCND1-*manipulated cells (**Figure 2F, S2B, Supplementary Table 2**). Together, these results demonstrate that the cell cycle has a very small effect on proteome composition and therefore does not confound the size-dependent proteome remodeling observed in large cells.

### Protein turnover plays a minor role in determining the proteome’s size dependence

One way in which a protein’s cellular concentration can change is through its turnover (**Figure 3A**). Protein turnover comprises combined effects of both proteolytic degradation and dilution by cell growth (*i.e.*, replacement with newly synthesized proteins). Previous studies have used metabolic labeling strategies to show that the turnover rates of individual proteins can vary widely.^52–55^ To test if protein turnover was size-dependent and contributed to proteome remodeling, we leveraged the semi-steady state nature of our *CCND1* overexpression/knockdown system to employ a pulse SILAC-TMT mass spectrometry approach (adapted from Zecha et al., 2018; see methods).^56^ Briefly, we first generated small and large populations of proliferating cells by culturing *P_TRE_ shCCND1* and *P_TRE_ CCND1* cells with doxycycline for five days. We then switched to media containing isotopically-labeled lysine and arginine to track the turnover of proteins over a ten day period. Individual timepoints were subsequently pooled together using isobaric TMT labels to measure SILAC ratios for the entire time course in a single multiplexed sample (**Figure 3B**).

**Figure 3.**
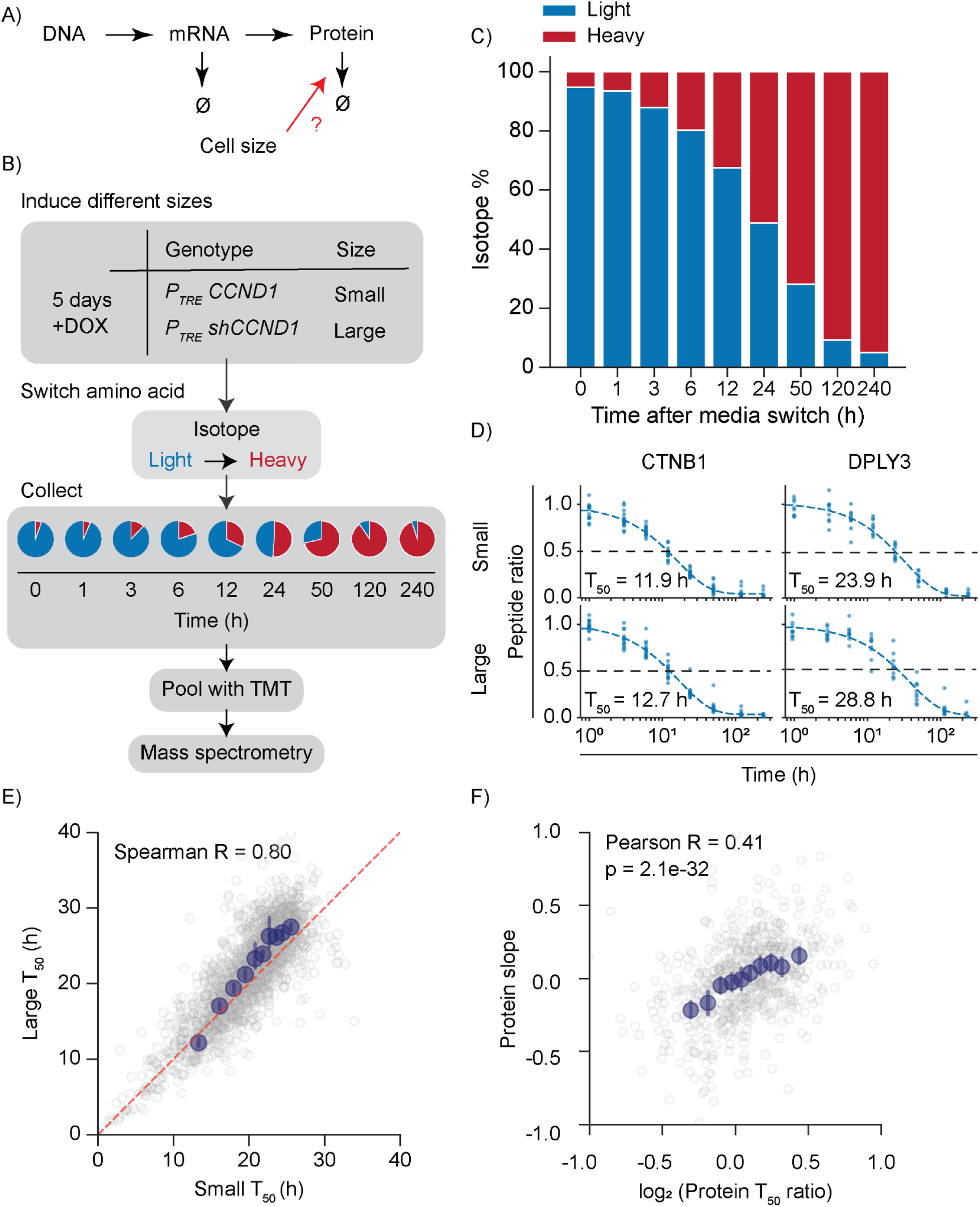
Protein turnover makes a minor contribution to size-dependent proteome remodeling. **A)** Schematic illustrating how size-dependent regulation of protein half-life could affect proteome composition. **B)** Outline of the pulse SILAC-TMT workflow used to globally measure the half-life of individual proteins in small and large cells. *P_TRE_ shCCND1* and *P_TRE_ CCND1* cells were induced with doxycycline for 5 days and allowed to grow to their small or large steady-state. Growth media was then swapped to media containing isotopically-labeled lysine and arginine for the next 10 days. Pie graphs illustrate the progressive intracellular incorporation of isotopically heavy amino acids into newly synthesized proteins. Timepoints were pooled together using isobaric TMT labels to measure SILAC ratios for all collected timepoints in a single multiplexed sample. For the biological replicate, isotope labels were reversed in pulse-chase order (**Supplementary Figure 3D, 3E**). **C)** Example bar plot showing the proportion of light and heavy labeled peptides at each timepoint in the time course. In this example, peptides labelled with lighter versions of isotopes were chased. **D)** Example decay curves of selected proteins in small and large cells. Decay curves were fitted using all peptide measurements for a given protein as previously described.^56^ The obtained decay rate was used to estimate protein half-life. **E)** Correlation of protein half-life (T_50_) measured from large and small cells’ protein turnover rates. The red dashed line marks the y=x line. Blue dots are the binned averages, and error bars represent the 95% confidence intervals. N = 2018 proteins. **F)** Correlation of size-dependent protein concentration changes with size-dependent changes in protein half-life. Only proteins that had half-lives shorter than the cell doubling time were examined. A protein’s T_50_ ratio is its half-life in large cells divided by that in the small cells. For clarity, x and y axes ranges are limited from -1 to 1. N = 638 proteins.

After establishing our labeling method, we determined the extent of the size-dependence of protein turnover and whether or not this contributed to size scaling. As expected, following the isotope switch, we observed the gradual replacement of one set of isotopically labelled proteins for another (**Figure 3C**). For individual proteins, we estimated their half-life (time to 50% replacement, or T_50_) by fitting a decay curve against all detected peptides mapping to their respective proteins (**Figure 3D, Supplementary Table 3**). At steady state, the rate of incorporation is theoretically equal to the rate of turnover. For simplicity, we opted to use protein half-life derived from turnover rates only, but we found that the protein half-life obtained from either rate was highly correlated (**Figure S3A**). The half-life obtained from our measurement also correlated well with a previously published study in *HeLa* cells, which used similar methods (**Figure S3B, S3C**).^56^ Overall, protein half-life was mostly constant across a near twofold size increase, indicating that the turnover rate of proteins are independent of cell size increase (**Figure 3E, Supplementary Table 3**).

It has previously been shown that in cycling human cells, the majority of proteins are regulated at the level of dilution rather than degradation.^56^ For a protein concentration to be primarily regulated by degradation, its half-life would need to be shorter than the time scale associated with dilution, which is the time scale of growth and division. In other words, proteins would need to be degraded more quickly than the average cell cycle duration. We identified these short-lived proteins by estimating the cell cycle doubling time of our small and large cells through microscopy (see methods). On average, our population of small cells (*P_TRE_ CCND1* + doxycycline) divided in ∼19 hours, while the large cells (*P_TRE_ shCCND1* + doxycycline) divided in ∼25 hours. Isolating our analysis to these short-lived proteins, when we compared the changes in protein turnover rate against the proteome size slope, we found a small, but significant correlation that partially explains the change in size-dependent proteome remodeling (**Figure 3F**). Therefore, protein turnover plays a significant, but minor role in shaping the protein composition of cells for short-lived proteins as they increase in size.

### Size-dependent transcriptomic remodeling largely underlies proteome remodeling

If post-translational regulation plays only a minor role in size-dependent proteome remodeling, then transcriptional regulation may explain a majority of the protein-level changes (**Figure 4A**). In this model, we should expect that the fold changes observed for protein concentrations correlate with those observed at the mRNA level. Protein changes are, in general, known to be correlated with changes at the mRNA level, with typical Pearson R correlations between mRNA and protein levels around 0.6.^57–59^ In our previous study using budding yeast, cells arrested for varying time to generate differently sized cells showed high correlation between fold changes in mRNA with those of proteins with a Pearson R value of 0.63.^33^ In contrast, our original study of differential protein scaling in cultured human cells found only a moderate correlation between the size-dependence of mRNA and protein concentrations (Pearson R value of 0.29).^30^ This discrepancy may have arisen from the long sorting process to isolate cells of different sizes used in the original study, which could confound transcriptomic measurements because of the relatively short half-life of mRNA on the time scale of hours.

**Figure 4.**
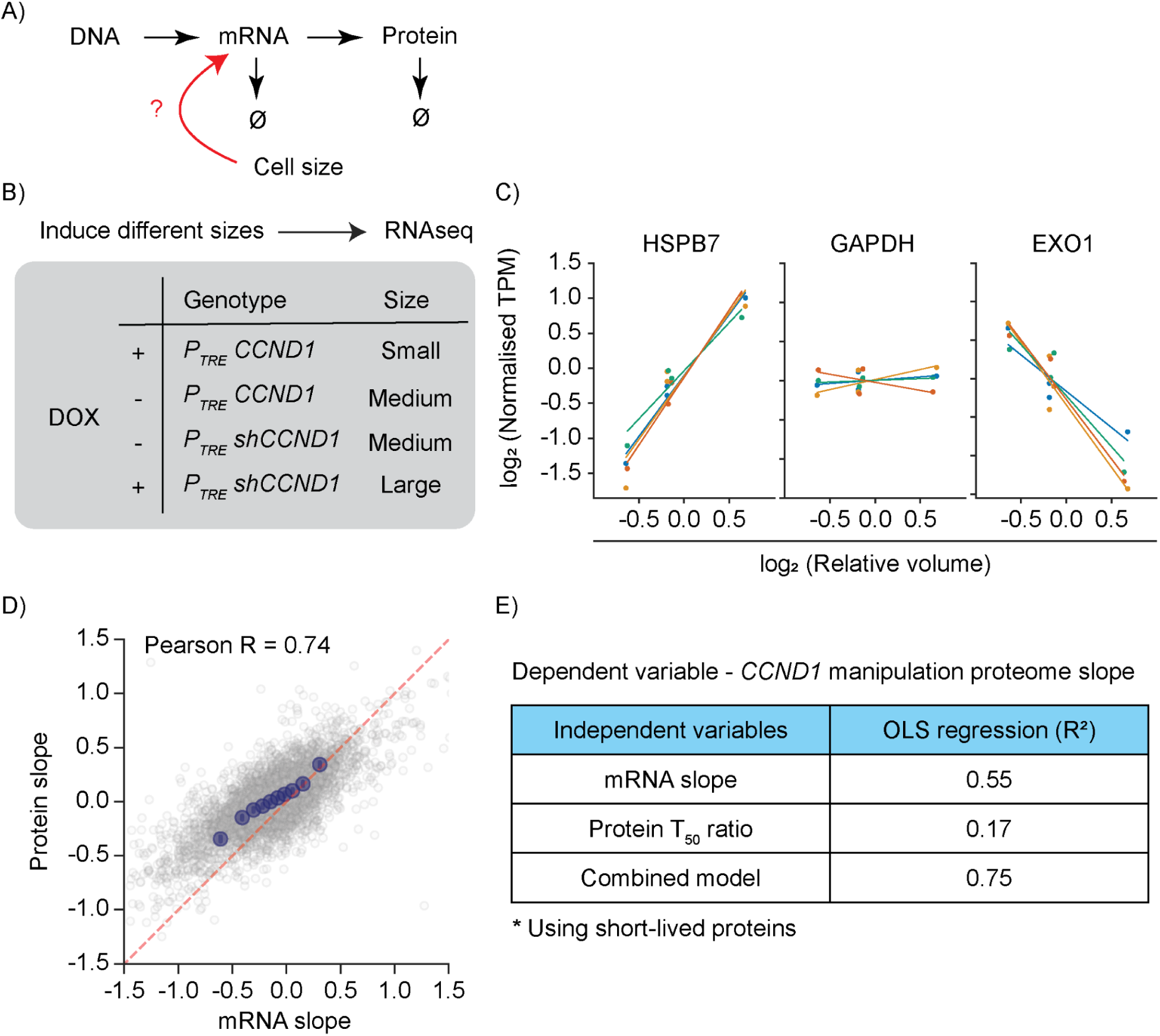
Transcriptome size-dependence explains most of the proteome remodeling. **A)** Schematic illustrating how size-dependent transcriptional regulation could affect proteome composition. **B)** Outline of RNAseq experiment to measure transcriptomic changes with cell size. *P_TRE_ shCCND1* and *P_TRE_ CCND1* cells were grown to their target sizes with doxycycline induction or DMSO for 4 or 5 days before collection. **C)** Examples of super-scaling, scaling, and subscaling mRNAs. Individual points mark the transcript reads mapping to labelled genes, and the colors denote different biological replicates. Linear regression lines through points in individual replicates are shown. Slope values for mRNAs were taken as the average of all individual replicates’ slopes. N = 4 replicates. **D)** Correlation of protein slopes with mRNA slopes. mRNA slopes were calculated as described in Figure 1B using the relative change in TPM. The red dashed line marks the y=x line. Blue dots are binned averages. N = 4720 genes. **E)** Ordinary Least Squares regression model to predict size-dependent protein concentration changes using protein half-life ratios and mRNA slopes as input variables. The model uses genes common to both datasets and the half-life of short-lived proteins (see methods).

To avoid the potential artifacts due to cell sorting, we used our *CCND1* induction system to quickly process differently-sized cell populations for RNAseq analysis (**Figure 4B**). As before, we extracted the relative change in transcript concentration with increasing cell size (**Figure 4C, Supplementary Table 4**). In contrast to our previous study using sorted human cells, this quicker method of extraction yielded highly-correlated mRNA fold changes with protein fold changes (**Figure 4D, Supplementary Table 4**). Combining both transcriptomic and protein half-life measurements, we were able to predict protein concentration changes with R^2^ = 0.75, with most of the explanatory power coming from the transcriptomic information (**Figure 4E**). Therefore, size-dependent protein remodeling appears to mostly originate at the level of transcription.

### Size-dependent mRNA turnover does not explain transcriptomic remodeling

The composition of the transcriptome is determined by both synthesis and degradation. In our previous study in budding yeast, we found that bulk mRNA was stabilized with increased cell size to maintain mRNA concentration homeostasis.^60^ However, we did not test whether there were any differential changes in mRNA stability that could contribute to the size-dependence of mRNA concentrations (**Figure 5A**). We therefore tested if mRNA turnover is important for remodeling the transcriptome with increasing cell size.

**Figure 5.**
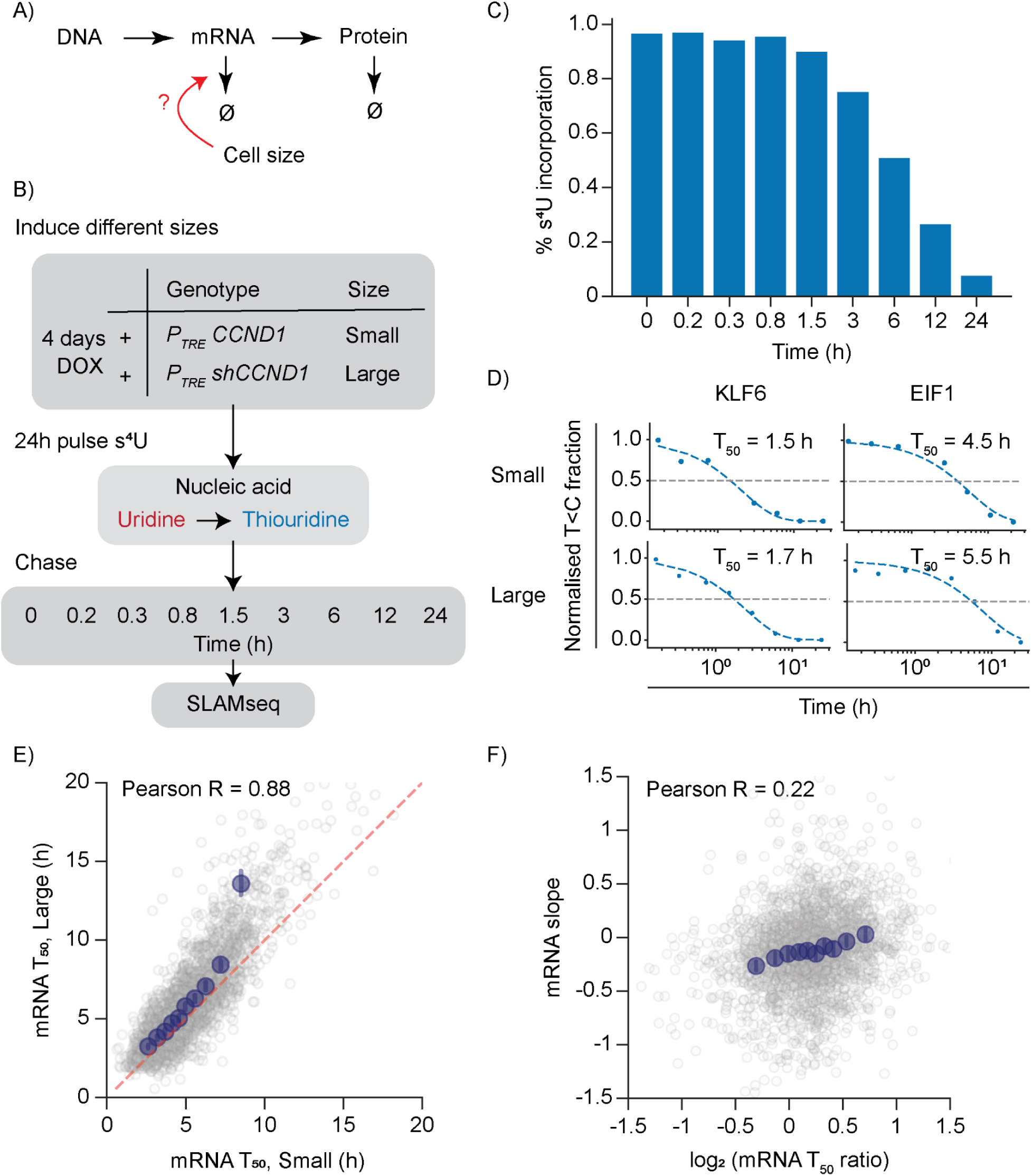
mRNA half-life of genes do not differentially change with increased cell size. **A)** Schematic illustrating how size-dependent regulation of mRNA half-life could affect mRNA concentrations and thereby affect proteome composition. **B)** Experimental setup for size-dependent SLAMseq. *P_TRE_ shCCND1* and *P_TRE_ CCND1* cells were induced to new sizes for 4 days, then pulsed for 24 hours with thiouridine. The pulsing medium was replaced every 3 hours, and doxycycline was maintained throughout the entire pulse and chase periods. After the pulse period, the medium was replaced with excess uridine, and samples chased at various timepoints for 24 hours before collection for SLAMseq. **C)** Percentage incorporation of thiouridine in mRNA as measured by T to C substitution fraction induced during iodoacetamide treatment and alkylation in one example sample. **D)** Example fit decay curves of mRNA for different cell sizes. Individual points mark the T to C substitution fraction for the given gene at that time. Data are normalized to the first time point (0 hour). Decay rates extracted from fitting were subsequently used to calculate mRNA half-life (see methods). **E)** mRNA half-life of genes do not show differential changes with increase in cell size. Red dashed line marks the y=x line. Blue dots indicate the binned averages, and error bars the 95% confidence intervals. N = 2536 genes. **F)** Correlation of size-dependent changes in mRNA half-life with size-dependent changes of mRNA concentrations. mRNA’s T_50_ ratio is its half-life in large cells divided by that in the small cells. N = 2437 genes.

To test if mRNA turnover contributes to transcriptome remodeling with cell size, we employed SLAMseq, a metabolic RNA labeling approach that involves pulsing of the media with a uridine analog, thiouridine (s^4^U). This analog can then be converted to cytidine through an alkylation reaction and subsequently detected by mRNA sequencing as a T to C nucleotide substitution.^61^ During the chase period, the mRNA incorporating thiouridine gradually decays, which enables the mRNA half-life measurement. To test for size-dependence, we used our genetic system to generate populations of differently sized cells on which we performed SLAMseq. Briefly, we pulsed the media with thiouridine for 24 hours and then replaced the media with excess uridine to minimize further incorporation of thiouridine (see methods). We then collected samples over the next 24 hours for SLAMseq analysis (**Figure 5B**). By following the proportion of T to C substitution introduced by the incorporation of thiouridine, we could monitor the gradual decay of mRNA species bearing the nucleoside analog over time (**Figure 5C**). We fit the decay curve for each mRNA for respective cell sizes to determine the mRNA half-life (**Figure 5D, Supplementary Table 5**). The median mRNA half-life of all genes for medium sized cells was 6.7 hours, which is in agreement with previous estimate of ∼6.9 hours for *HeLa* cells.^62^ Moreover, the mRNA half-lives of individual genes correlated well with those measured in a previous SLAMseq study of mESCs (**Figure S5A**).^61^ To determine the size-dependence of the turnover of mRNA transcripts, we then obtained the mRNA half-life ratio by dividing the mRNA half-life in large cells by that in small cells. We observed that all genes, except for those belonging to the top ∼10% in mRNA half-life, did not show marked differential changes in half-life with increase in cell size. Instead, on average, the mRNAs showed a small absolute increase in half-life (**Figure 5E, Supplementary Table 5**). This is consistent with our previous work showing a broad stabilization of mRNA in larger yeast cells, which functioned to maintain mRNA concentrations in larger cells by counter-balancing the sub-scaling of mRNA synthesis.^60^ An analysis of our data revealed only a weak correlation between mRNA slope and mRNA half-life ratio, indicating that differential size-dependent effects on mRNA half-life do not underlie transcriptome remodeling (**Figure 5F**). We therefore conclude it is likely that size-dependent changes in transcription are mostly responsible for size-dependent transcriptome, and thereby, proteome remodeling.

### Size-scaling transcription is driven by an increase in burst length

Knowing that mRNA synthesis is key to understanding how mRNA composition differentially scales with cell size, we first sought to understand how mRNA scaling may be achieved through synthesis. Transcription occurs in bursts, where periods of dormancy are punctuated by periods of intense transcriptional activity so that the promoter can be seen to exist in at least two distinct states.^63–66^ These transitions are controlled in the promoter region, which serves as the nexus of many different transcriptional regulators, including chromatin modifications, nucleosome occupancy, enhancer-promoter contacts, and binding of transcriptional factors. All such modes of transcriptional regulation culminate in several kinetic bursting parameters, including burst size, amplitude, and ON/OFF times. These parameters are related to the number of transcripts produced during a productive burst period, average loading rate of RNA polymerase II, and the productive/inactive period of transcription respectively. Previous work has shown that transcript abundance scales with the increase in cell size in both prokaryotes and eukaryotes.^60,67–70^ In fixed human cells, smFISH data measuring the distributions of mRNA counts was used with a mathematical model of expected variability to infer that an increase in burst size primarily drives the scaling of transcription with cell size.^70^ However, there has been no real-time analysis of transcriptional bursts to confirm the scaling of burst size with cell size, and it is unclear whether there are other factors contributing to transcriptomic scaling.

To determine how transcriptional burst parameters are modulated by increasing cell size, we sought to visualize transcription in real-time using the MS2 system to fluorescently label individual mRNA molecules at the site of transcription.^71–75^ We performed this size-dependent burst analysis using near-diploid *HBEC-kt3* cells that have MS2 stem loops incorporated into their genes’ endogenous introns (**Figure 6A**).^76^ We chose to image the bursts of two genes, *RAB7A* and *RHOA*, which we classified to be scaling in our *RPE-1* RNAseq data **(Supplementary Figure 6A, 6B)**. After imaging these cells in their natural cycling state, we segmented and tracked their burst dynamics using a custom algorithm (**Figure 6B**) (see methods). ON and OFF states were determined based on intensity thresholding. Since we did not have a cell cycle marker, we instead segregated cells by their nuclear area, and took a range of cells within a nuclear size window that we knew to be highly enriched in G1 (**Supplementary Figure 6C, 6D**). From our traces, we extracted features of transcriptional bursts and asked how they changed with increasing nuclear volume in G1 (**Figure 6C**). We used nuclear area measurements to estimate the nuclear volume, which we previously found to be highly correlated with cell size.^77^ In agreement with previous work,^67,70^ we found that both *RAB7A* and *RHOA* scaled their burst size with cell size, indicating increased number of RNA polymerase II loading onto the promoter during initiation, or the increase in number of RNA polymerase II released in a given pause-release. Decomposing burst size into ON time and average burst amplitude, we found that the scaling of burst size originates mainly from the scaling of ON time, meaning that once the gene is activated, it remains activated for a longer time in larger cells than in smaller cells. Burst amplitude remained relatively constant. Inactive periods between productive transcription are also shortened, although the magnitude of this change was not as large as the change in ON time (**Figure 6D, Supplementary Figure 6E, Supplementary Table 6, 7**). Taken together, our data indicate that as cells get larger, they scale up transcription mainly by prolonging the productive burst period but also, to a lesser extent, by more quickly reactivating transcription.

**Figure 6.**
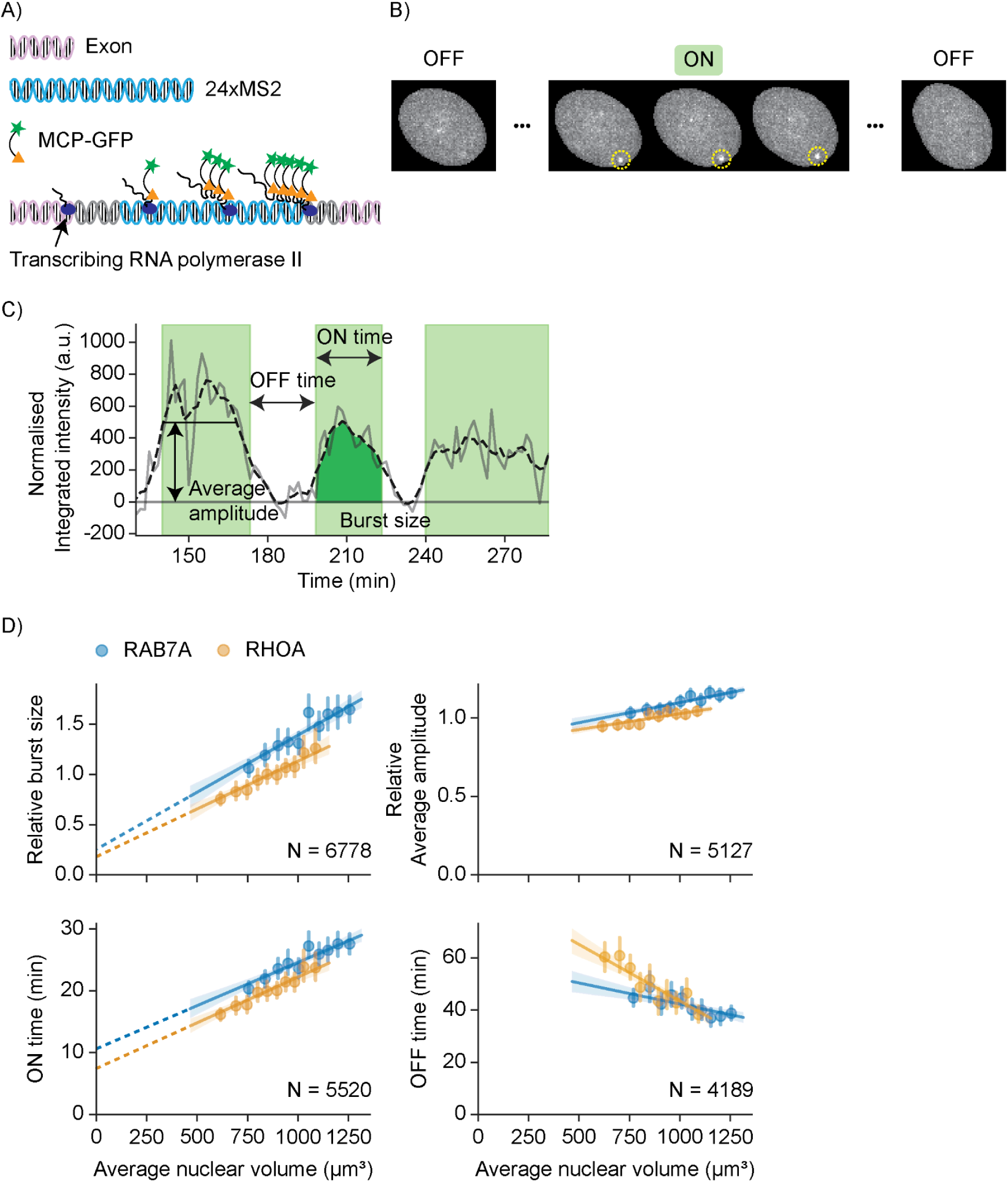
ON time increase scales with increase in cell size. **A)** MS2 stem-loop knock-in to endogenous introns of genes to visualize real-time transcription. Transcripts become visible when GFP-conjugated MCP proteins bind to transcribed MS2 stem loops. Cell lines were from a previously published study.^76^ **B)** Example segmentation and tracking of bursts. Bright spots in individual nuclei were classified, segmented, and tracked over time (see methods). **C)** Example trace of a transcriptional burst at a single locus. Transcriptional burst parameters of interest are indicated. ON time is the length of the productive bursting period. OFF time is the length of inactive period between productive bursts. Burst size is the area under the trace curve and is related to the total number of transcripts produced in each burst. Average amplitude is the burst size divided by ON time and relates to the average loading rate of RNA polymerase II during initiation. **D)** Burst parameters change with increases in cell size. Individual points mark binned averages, and error bars mark the standard error of means. Dashed lines denote the extended line of best fit that crosses the y-axis near the origin. Here, a line that crosses the origin is indicative of perfect scaling. For *RAB7A*, 6778 active transcriptional periods from 1943 cells were analyzed, and for *RHOA*, 5127 periods from 1296 cells were analyzed.

## Discussion

Understanding how cell size influences cell physiology is essential for understanding why cell size is tightly regulated and different types of cells adopt different characteristic sizes. Cell physiology is most holistically represented in its proteome composition, which changes with size. Such a size-dependent proteome remodeling is highly conserved as it has now been measured in multiple different human cell types using orthogonal experimental strategies, as well as in both yeast and *E. coli*.^30,31,33,34^ While the work in these unicellular organisms suggested that transcriptomic changes account for most of the size-dependent changes in protein composition, it was not known if this was also true in human cells, and whether post-transcriptional and post-translational regulation also contribute significantly as suggested by our earlier study.^30^

To identify the mechanistic origin of size-dependent proteome remodeling in human cells, we generated cell lines cycling at different sizes to measure protein and mRNA concentrations and their turnover rates. Our analysis revealed that differential size-dependent transcription drives the size-dependence of proteome remodeling in human cells. For two genes whose transcription scales in proportion to cell size, we show that this mostly results from an increase in the duration of a transcriptional burst and that changes in the duration of promoter inactivity and transcription amplitude were more minor (**Figure 7A**). Size-dependent proteome remodeling is primarily driven by the differential scaling of transcription at individual genes, whereas mRNA and protein turnover rates are largely robust to changes in cell size and thus only weakly contribute to proteome remodeling (**Figure 7B**). Overall, our findings provide the most detailed investigation of size-dependent gene expression pathways to date.

**Figure 7.**
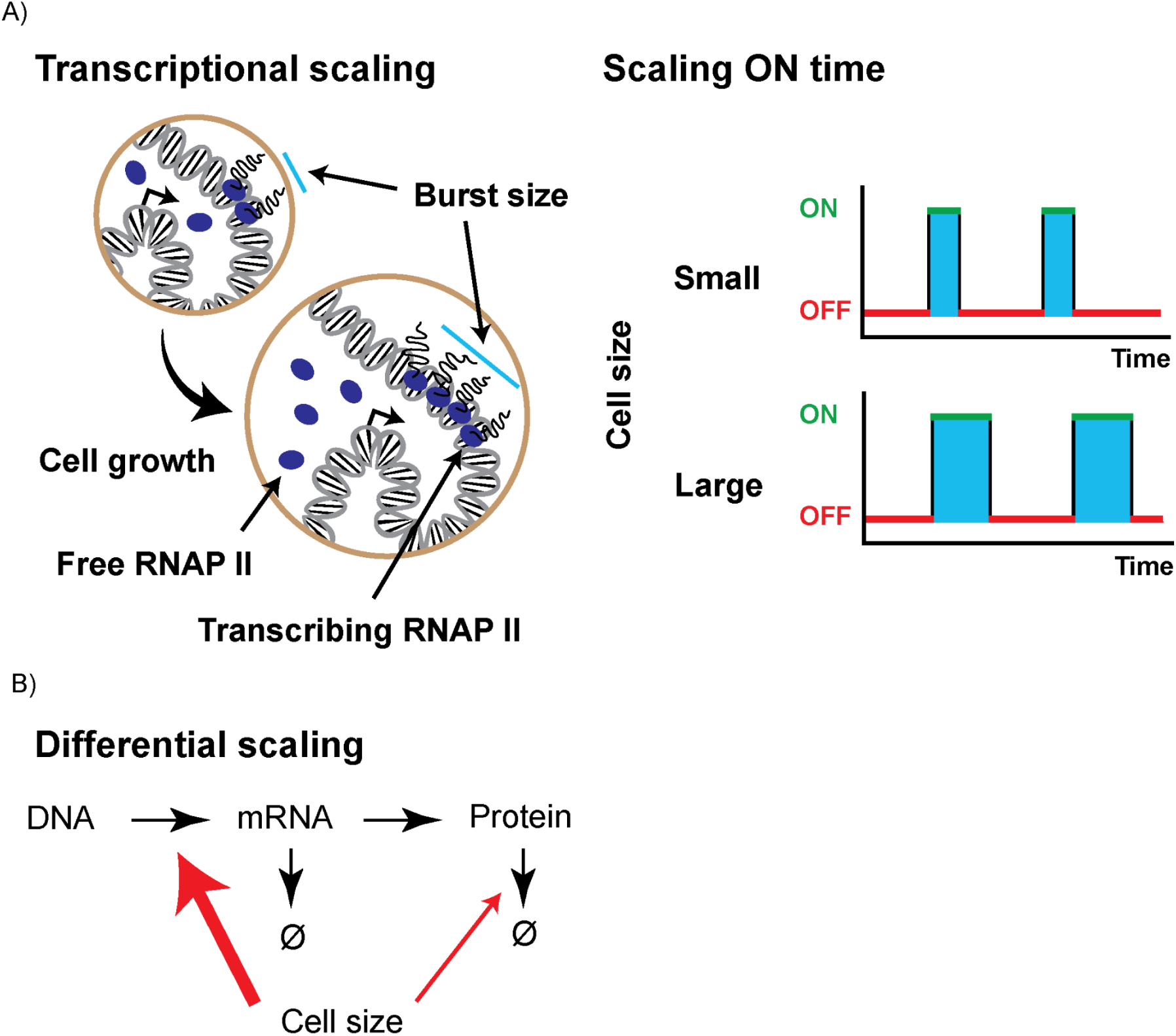
Size-dependent transcription underlies proteome remodeling with cell size. **A)** To maintain scaling transcription with increased cell size, cells keep the promoter productive for longer for RNA polymerase II (RNAP II) binding once turned on. Burst size is the total number of transcripts produced in a single burst. Changes to OFF time and average burst amplitude are relatively small and not depicted. **B)** Summary of cell-size dependent changes that occur along the gene expression pathway. Arrow sizes indicate strength of contribution of process to size-dependent proteomic remodeling. For differential protein scaling, transcription is the highest contributor, with a modest addition from protein turnover. mRNA turnover is least affected by the increase in cell size to achieve differential protein scaling. Effects of translation were not examined in this study.

### Size-dependent transcriptional bursting scales gene expression

After identifying a primarily transcriptional basis for size-dependent proteome remodeling, we sought to mechanistically probe how increasing cell size drives changes to specific features of transcription to achieve uniform scaling. Specifically, we examined rate parameters related to transcriptional bursting dynamics, namely their rates of initiation, duration, and amplitude.^64,78^ To see how cell size modulated these features of transcription, we used live-imaging of transcriptional loci to examine two endogenous size-scaling genes, *RAB7A* and *RHOA*. The transcription of these two genes scales in proportion to cell size predominantly due to longer ON times, but also from contributions from the initiation rate and amplitude. This is consistent with previous smFISH studies concluding that the total amount of transcripts in a burst increases with cell size in human cells.^67,70^ However, these previous studies lacked the resolution to further decompose the relationship between transcriptional bursts and cell size because they lacked the detailed temporal information of burst dynamics that we provide from our live-cell analysis.

The increase in burst size driven by increased ON times of *RAB7A* and *RHOA* echoes previous studies linking *MYC* with modulation of burst size and ON time.^79,80^ *MYC* over-expression drove a global upregulation of transcription through an increase in burst size mainly due to an increase in ON length.^79^ Single molecule tracking of core transcriptional regulators such as *TBP*, *SPT5*, and *MED1* showed a change in their dwell time, which may explain the increase in ON length. While it is unclear whether *MYC*, a well-known growth regulator, is responsible for the size-dependent modulation of transcriptional burst parameters, we speculate that such a mechanism could be at play through various activating transcription factors. For example, larger cells may have a greater number of transcription factors, but a constant number of binding sites on the genome. This leads to an effective increase in transcription factor binding and promoter occupancy that can increase transcription in large cells similar to *MYC* overexpression. Future studies could test this by examining residence times of transcription factors at the promoters of scaling genes using single molecule microscopy methods.^81^

Pause-release is also another potential point of regulation, especially since the release of paused RNA polymerase II is thought to be limiting for human transcription.^82–84^ Changing activities of positive and negative regulators of transcriptional pause-release, such as p-TEFb, NELF, or DSIF, could act to prolong the ON time. Future identification of the transcriptional burst modulators that change activity with cell size will be essential in understanding how cells integrate growth signals to increase their transcriptional output.

### What is the molecular basis for size-dependent transcriptional remodeling?

Though the two model genes whose bursting parameters we measured illustrate how transcriptional output scales in proportion to cell volume, this proportional scaling clearly does not occur at all genes. Indeed, individual genes can couple the scaling of their transcription to cell volume to varying degrees and thereby drive the differential transcriptome remodeling we and others have observed.^31,33^

A recent study investigating how the transcriptome changes with cell size in budding yeast proposed that size-dependent changes to the transcriptome are driven by transcriptomic “inversions”.^85^ Similar findings of size-dependent scaling of transcription based on basal expression were reported for genome-diluted intestinal cells in *C. elegans*.^86^ These inversions occur when RNA polymerase binding saturates at highly expressed genes and therefore results in their relative down-regulation in large cells. If this model were true for mammalian cells, we would expect to see bias in our mRNA slopes measurements, with mRNAs with higher basal expression levels having more negative slopes. However, we do not see any correlation between the basal expression level of an mRNA and its propensity to super- or sub-scale with size (**Supplementary Figure 4A**).

We speculate two possible explanations for how differential scaling could occur. First, each individual gene is encoded in a way that predetermines how the binding and bursting of transcriptional machinery will scale with volume. While the two example genes we measure increase transcription linearly with volume, other genes could be encoded to adopt different trajectories and thus result in differential scaling. Though it remains unclear how this gene feature would be encoded, it would present interesting evolutionary implications, since the decision to tune up or down the transcription of certain genes in large cells would be hardwired in the genome. Alternatively, differential scaling could be explained by an upstream regulatory signal that gradually changes as cells grow larger and thereby alters the transcriptional landscape. This possibility is supported by the fact that size-dependent transcriptional remodeling closely resembles the transcriptional changes associated with mTOR inhibition.^33,35^ Finally, the size-dependent proteome remodeling is dependent on the dilution of the genome with increasing cell size.^30,31,33,43^ Thus, changes in growth-related signaling that coincide with dilution of the genome could represent an upstream signal that drives the differential scaling of the transcriptome.

### mRNA and protein turnover do not show differential changes with cell size

As proliferating cells grow larger, proteome changes typically associated with cellular senescence are increasingly pronounced,^30^ and when cells become too large, they are no longer able to re-enter the cell cycle and therefore become functionally senescent.^37,39–42^ Since senescence has been linked to broad defects in proteostasis,^87^ we hypothesized that protein and mRNA turnover may be affected by increases in cell size. However, the size-dependence of both protein and mRNA half-lives are largely unchanged in the larger proliferating cells we measured here. We do observe a small population-wide increase in the mRNA half-life with increase in size (**Figure 5E**), which is consistent with the previous observation that yeast mRNA is stabilized as cell size increases to maintain a constant total mRNA concentration.^60^

A recent study examined the relationship between cell size and protein degradation rates and reported that large cells hyper-activate their proteasomes to upregulate protein degradation.^88^ The authors found that increasing cell size by blocking cell cycle progression and, in conjunction, also acutely inhibiting the proteasome yielded greater accumulation of polyubiquitinated lysine (K48-polyUb), a major targeting signal for proteasomal degradation. From this, they concluded that large cells increase protein degradation. Blockade of CDK4/6 activity, however, which is analogous to our knockdown of *CCND1*, did not induce a similar accumulation of polyubiquitinated lysine. The authors attributed this difference to the lack of growth slowdown from CDK4/6 inhibition, which may be consistent with our finding that protein turnover rates generally do not change with increasing cell size. We also note that our study was done under more steady state conditions with respect to cell size supporting our conclusion that there is limited size-dependent protein degradation shaping the proteome’s size-dependence.

### Altered cell vulnerability with cell size deviation

Proteome “imbalance” is a central concern in overgrown cells with diluted genomes.^37,89–91^ Lysosomal proteins are among the highest super-scaling proteins in our dataset, and aberrant lysosomal function has been linked to various cell defects including senescence.^49–51^ We therefore reasoned that even cells close to their natural size range may be beginning to show altered lysosomal states. Indeed, we find that large proliferating cells have higher vulnerability to lysosomal perturbation by inhibition from both LLoMe and chloroquine. While our effect size was not large, in contexts where cells are much enlarged, for example following CDK4/6 inhibition, these lysosomal vulnerabilities may be a potential useful exploit for treatment.^92^

Finally, the cell-size dependent changes in biosynthesis that create unique vulnerabilities have also been reported in other contexts, with upregulation of *IRF4* following CDK4/6 inhibition sensitizing multiple myeloma cells to bortezomib, and increased dependence of large cells on *GPX4* from higher lipid and iron content making them vulnerable to ferroptosis.^93,94^ We speculate that there may be more unique cell size-dependent vulnerabilities stemming from the imbalance in biosynthesis that can be exploited for cancer therapy.

**Supplementary Figure 1.**
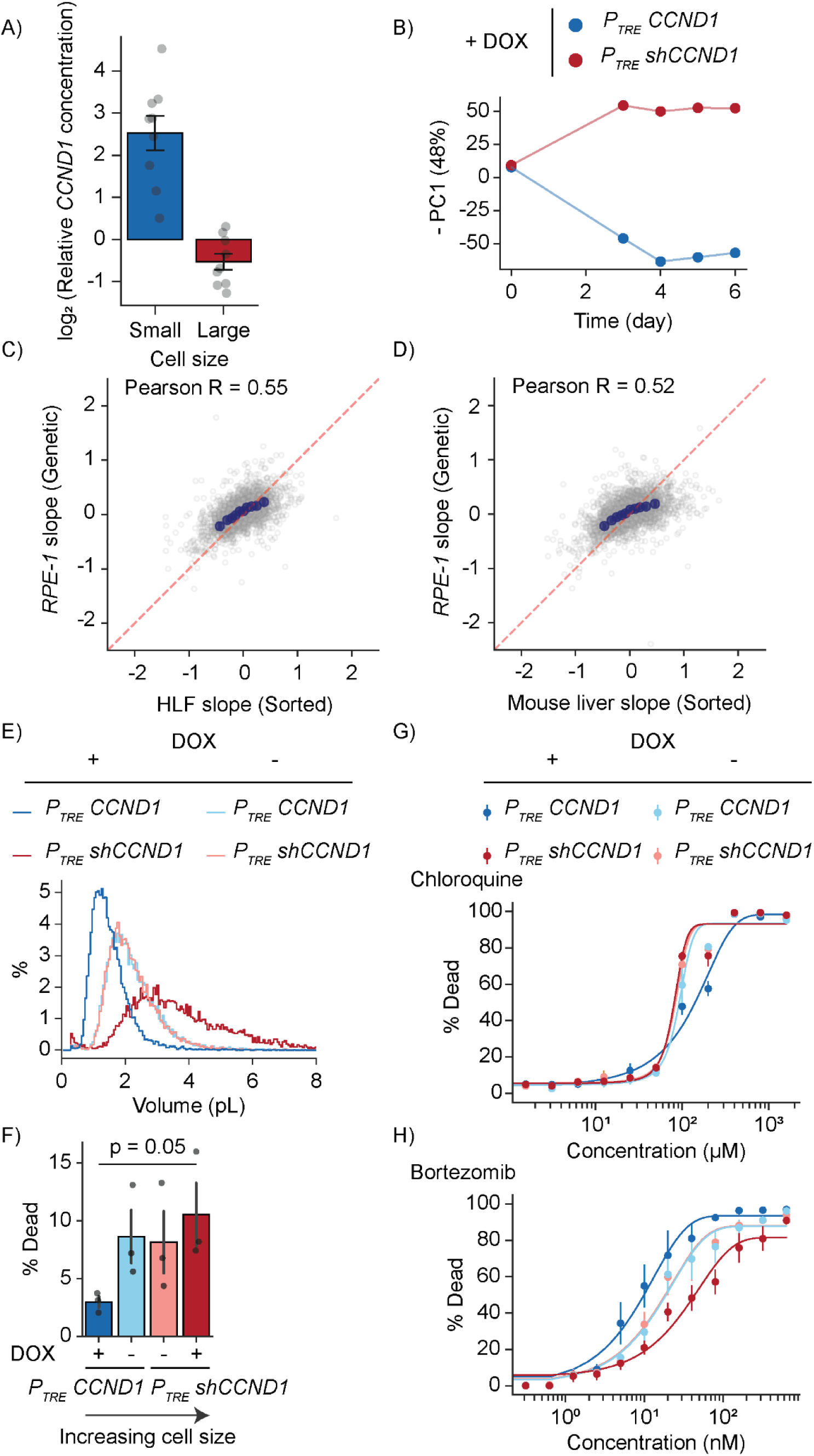
**A)** Relative *CCND1* peptide concentration as measured by mass-spectrometry following *CCND1* knockdown or overexpression with doxycycline induction. Data was normalized to DMSO treated cells before log2 transformation such that uninduced cells have an expression level = 0. Averages of 8 high-quality peptides were taken. Error bars mark the standard of the mean. **B)** Principal component analysis of *P_TRE_ shCCND1* and *P_TRE_ CCND1* cells’ proteomes as measured by TMT-mass spectrometry before and after doxycycline induction. Cells were collected on days 0, 3,4,5, and 6. Comparison of the first principal component against time shows the cells reaching proteomic steady state after the 4th day of induction. **C)** Correlation of protein slopes derived using the *CCND1* manipulation system in *RPE-1* cells with protein slopes previously measured using size-sorted primary human lung fibroblasts (*HLFs*) (N = 2560 proteins).^30^ **D)** Same as (C) but for mouse primary liver cells from a previously published study (N = 3127 proteins).^33^ **E)** Representative cell size distribution as measured by the Coulter counter for *P_TRE_ shCCND1* and *P_TRE_ CCND1* cells treated with doxycycline or DMSO for 5 days. **F)** Proportion of dying cells of the indicated conditions after 48 hour treatment of 50uM chloroquine, a lysosomotropic drug that perturbs lysosomal pH. Cell death was measured by Annexin V and a cell permeability dye. Baseline death rates from control (DMSO) conditions were subtracted to calculate death from chloroquine. P-value shows outcome of unpaired t-test. N = 2 replicates. **G)** Proportion of dead cells across a wide dosage range after 3 days of chloroquine. *P_TRE_ shCCND1* and *P_TRE_ CCND1* cells induced with doxycycline or DMSO to new sizes were treated with the drug and assayed for cell death using SYTOX DeepRed and microscopy. Quantification of cell death was carried out using a custom nuclear segmentation algorithm (see methods). N = 3 replicates. **H)** Same as (F), but for 2 days of treatment with bortezomib. N=3 replicates.

**Supplementary Figure 2.**
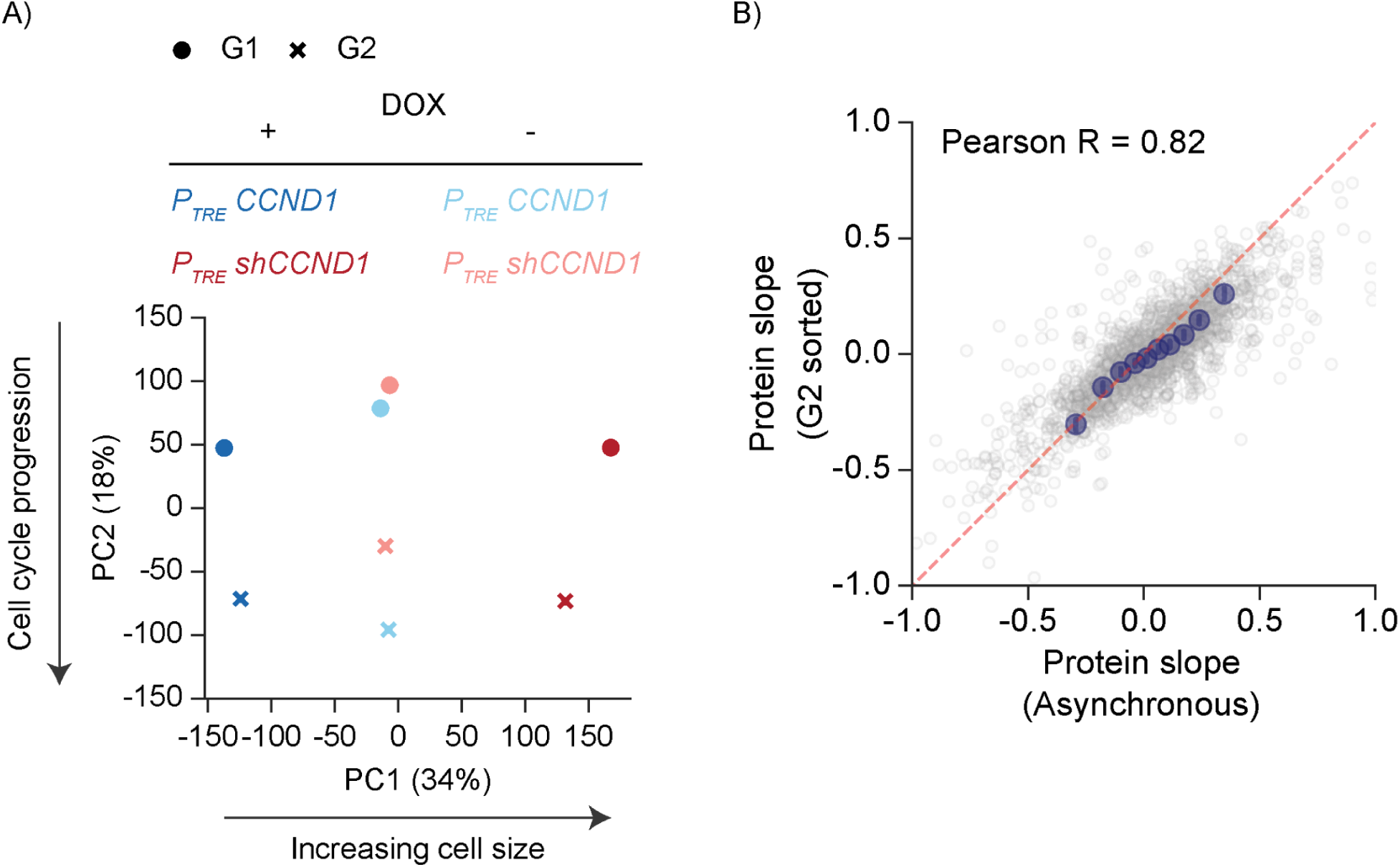
**A)** Principal component analysis of G1 and G2 sorted *P_TRE_ shCCND1* and *P_TRE_ CCND1* cells after 5 days of doxycycline or DMSO induction. Cells were sorted by cell cycle phase after DNA staining. Individual points show the proteome of indicated cells. Samples are separated by cell cycle and cell size, and samples in both cell cycle phases (G1 and G2) show the same pattern of change along PC1 (cell size). **B)** Protein slopes as obtained from asynchronous conditions (as in Figure 1F) correlated against protein slopes from cells sorted in G2. The two samples are highly correlated with one another, indicating that cell cycle effects do not play a major role in size-dependent proteome remodeling. The limited number of proteins (N = 1882 proteins) reflects the shallow depth of the proteome analysis for this control experiment.

**Supplementary Figure 3.**
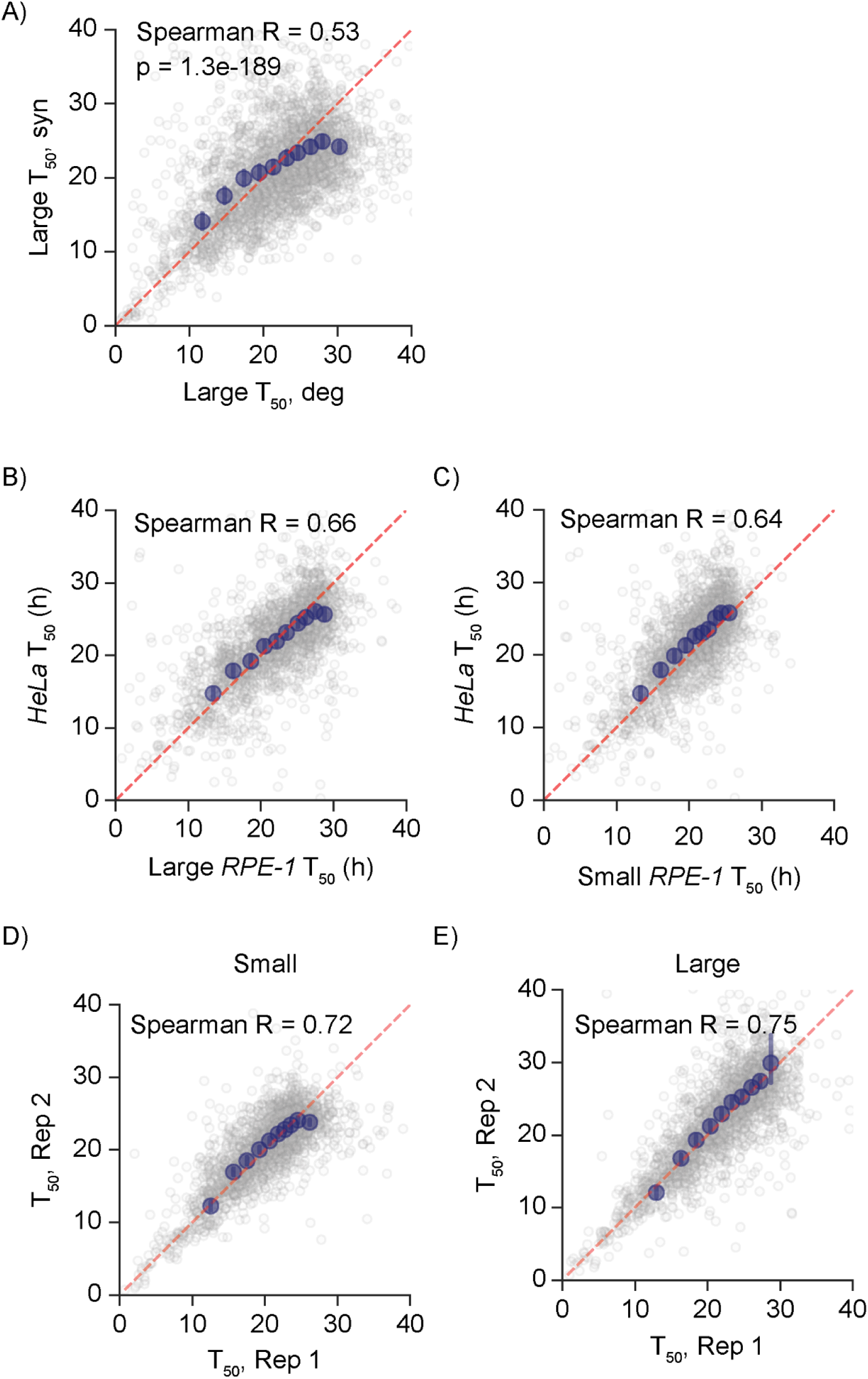
**A)** Protein half-life comparison as calculated from incorporation (Large T_50, syn_) and decay (Large T_50, deg_) rates for large cells. Red dashed line indicates the y = x line. Blue dots indicate binned averages, and error bars mark the 95% confidence interval. N = 2656 proteins. **B)** and **C)** Comparison of protein half-life obtained from our study (both small and large cells) against those from a previously published study in *HeLa* cells.^56^ All protein half-life values are from turnover rates. N = 1882 proteins for both plots. **D)** and **E)** Biological replicate comparison of protein turnover derived from small and large cells. N = 2018 proteins for both plots.

**Supplementary Figure 4.**
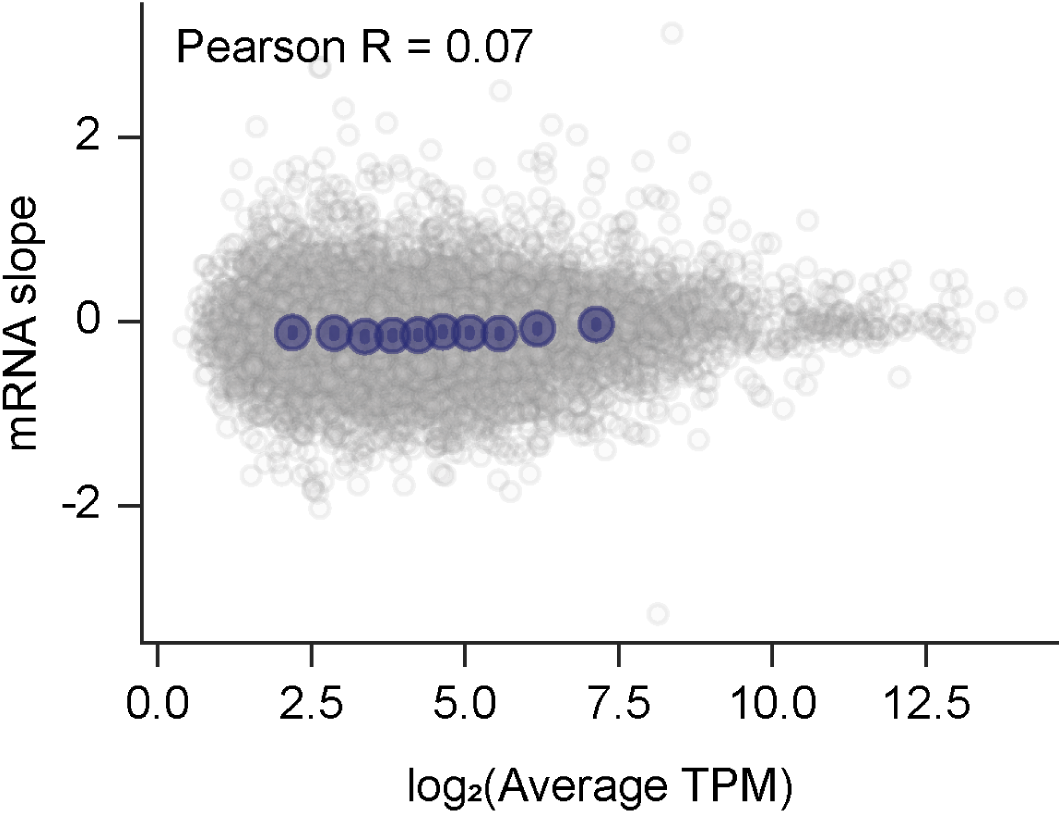
Size-dependent changes in mRNA concentration do not correlate with absolute expression levels. mRNA slope values were compared against the genes’ respective basal TPM values in medium-sized cells. Blue dots are binned averages. N = 10139 genes.

**Supplementary Figure 5.**
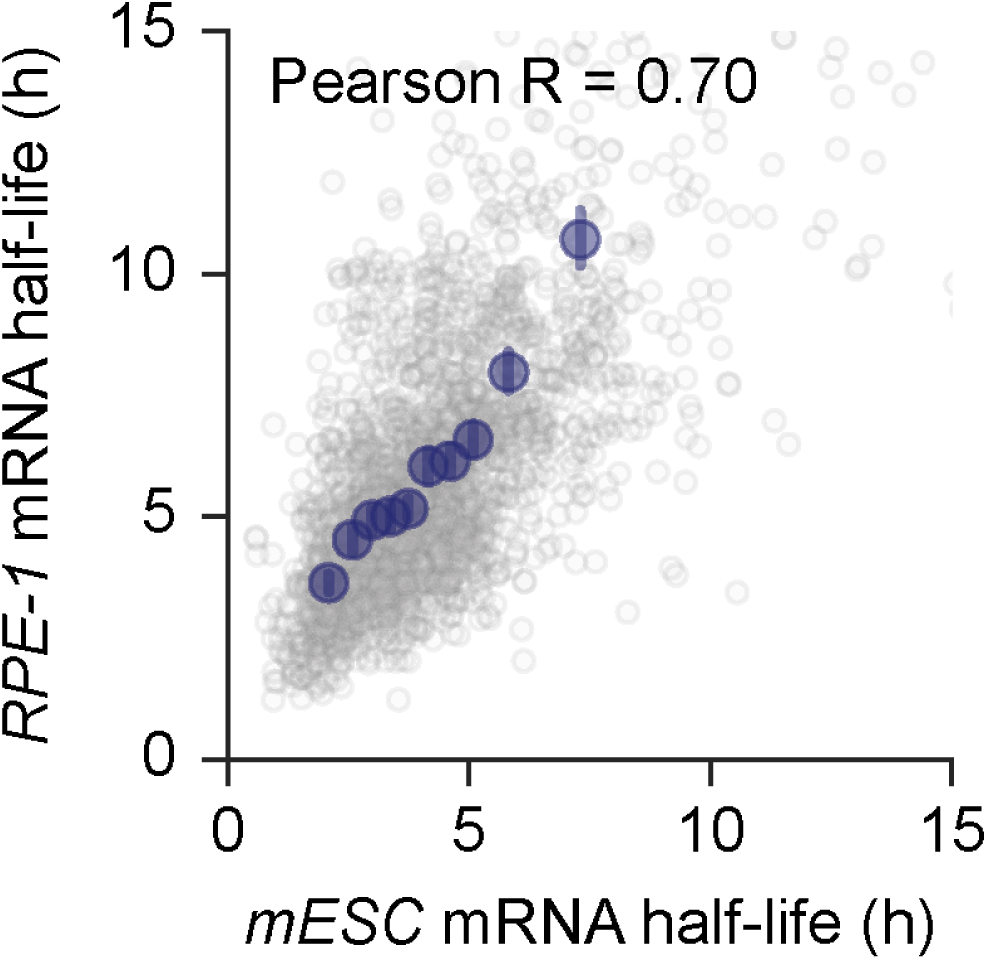
Comparison of mRNA half-life measured by this study and those from *mESC*s from a previously published study using SLAMseq.^61^ Average mRNA half-life of middle-sized cells were used for *RPE1* mRNA half-life. Mouse genes were humanized using the orthologs database from Mouse Genome Informatics. N = 2145 genes.

**Supplementary Figure 6.**
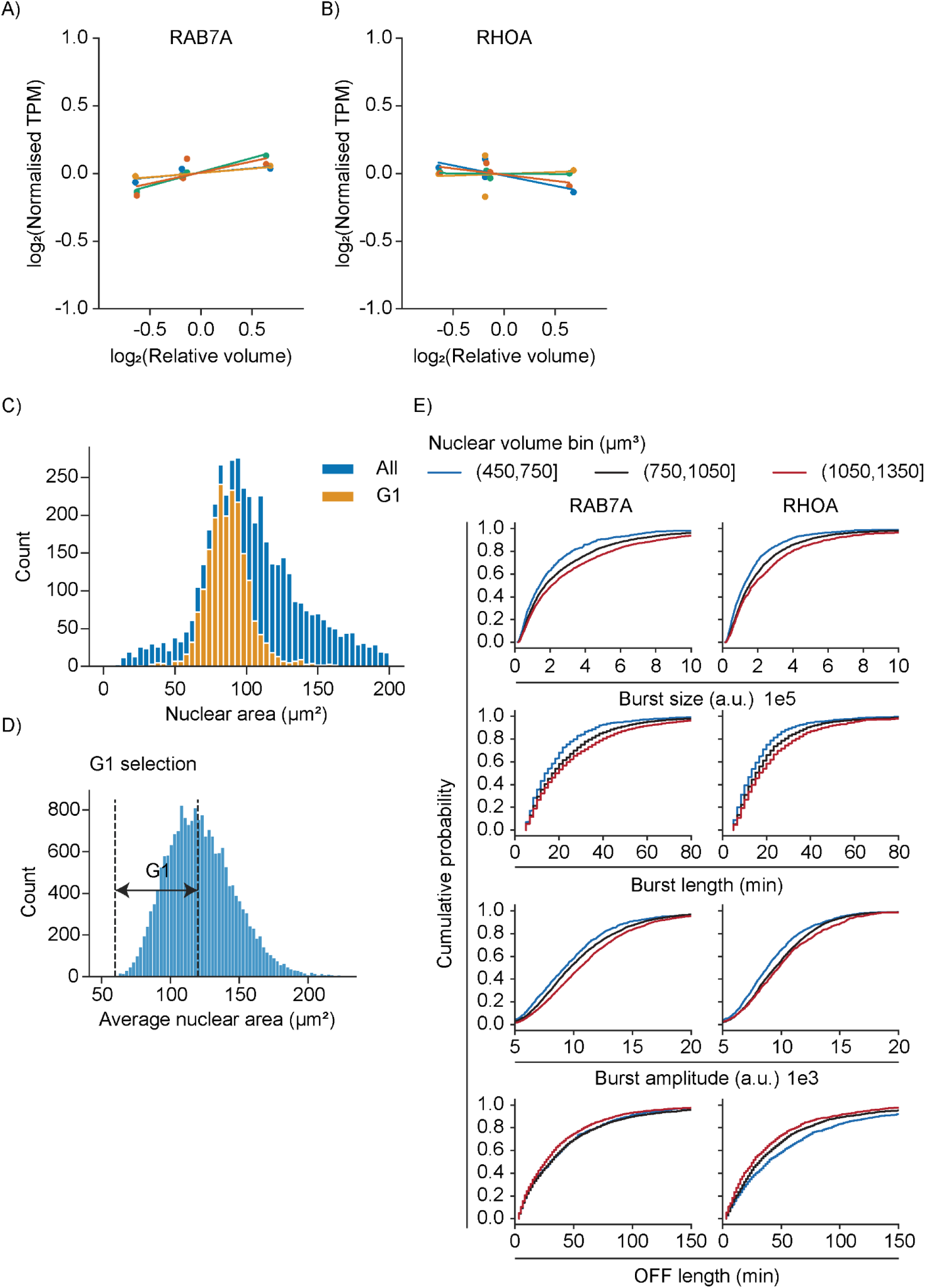
Size scaling mRNA concentrations for **A)** *RAB7A* and **B)** *RHOA* as measured by RNAseq. Individual points mark the transcript’s gene-specific mean-normalized TPM measured for a given size, and separate colors denote different replicates (N = 4 replicates). **C)** Nuclear area distribution of *HBEC-kt3* cells with endogenous *RAB7A* tagging of MS2 stem loops. G1 population distribution is highlighted to indicate G1 cells’ area distribution. Cells stained for DNA content with Hoechst were imaged with a widefield epifluorescence microscope separately from live-imaging experiments to determine nuclear size distribution of G1 cells. Nuclear area was used as a proxy for cell size. **D)** Window of selection of *HBEC-kt3* cells tagged with endogenous *RAB7A* tagging of MS2 stem loops for G1. These cells were selected for further downstream analysis of burst dynamics for this study. **E)** Cumulative distributive function plot of different burst parameters for genes *RAB7A* and *RHOA* binned by increasing cell size. Plots have been enlarged to highlight size-dependent changes in burst parameters.

## Methods

### Cell culture

Cells were cultured at 37℃ and 5% CO_2_ in Dulbecco’s Modified Eagle Medium/Nutrient Mixture F-12 (Corning; 10-092-CV) supplemented with 10% heat-inactivated fetal bovine serum (Corning; 35-011-CV) and 1% penicillin/streptomycin (Gibco; 15140-122). To induce cyclin D1 knockdown with or without wildtype cyclin D1 induction, 0.5 ug/mL doxycycline was added to the culture medium. Relatively steady states of cell size were reached by day 4 of doxycycline induction. MS2 stem loop incorporated human bronchial epithelial cells (*HBEC3-kt*) used in this study were from a previous study.^76^ Briefly, gene trap strategy was employed to integrate MS2 stem loops inside introns, after which the cells were single-cell isolated and genotyped to reveal their loci. *HBEC3-kt*s were grown in Airway Epithelial Cell Basal Medium (ATCC;PCS-300-030), supplemented with Bronchial Epithelial Cell Growth Kit (ATCC; PCS-300-040).

### Cyclin D1 replacement system in hTERT-RPE-1 cells

To establish our *CCND1* manipulation system in *RPE-1*s, the FLAG-tagged *CCND1* allele was PCR-amplified from the Rc/CMV-Cyclin D1 plasmid (kindly provided by Dr. Philip Hinds) and cloned into the pENTR/D-TOPO entry vector (Invitrogen, Cat. No. 45-0218) following the manufacturer’s instructions. *CCND1* wild-type was subsequently transferred into the pINDUCER20 lentiviral expression vector (Addgene; 44012) using the LR clonase reaction (Invitrogen, cat. No. 11791-020). An shRNA that targets the 3′-UTR of human *CCND1* mRNA (sequence: 5′-GCCAGGATGATAAGTTCCTTTC-3′) was cloned into the XhoI - MluI sites of the pINDUCER11 vector (Addgene; 44363), as previously described.^95^ Lentiviral particles were produced in 293T cells by co-transfection of either pINDUCER11 or pINDUCER20 with the packaging plasmid pCMV-dR8.2 dvpr (Addgene; 8455) and the envelope plasmid pCMV-VSV-G (Addgene; 8454), using X-tremeGENE 9 DNA transfection reagent (Sigma, Cat. No. 6365787001). Viral supernatants were used to transduce *hTERT-RPE-1* cells in the presence of 5 μg/mL polybrene (Sigma, Cat. No. 107689). Stable cell lines were selected by fluorescence-activated cell sorting (FACS) for GFP expression or by antibiotic selection with 400 μg/mL G-418 (GIBCO, Cat. No. 10131035), depending on the selectable marker encoded in the lentiviral vector.

### Cell size measurement

Cell sizes were measured by Z2 Coulter Counter (Beckman). Cells were first diluted in isoton solution, 0.5mL of the diluent taken for analysis, and the average of two measurements taken. Gates were set at 30.2 µm and 9 µm for upper and lower bounds respectively. For subsequent relative cell size calculations, the median cell size was taken.

### Doubling time measurement

Cells were induced with doxycycline for 4 or 5 days then seeded onto 35mm imaging dishes (MatTek P35G-1.5-20-C). After 2 and 3 days of growth, cells were fixed with 4% formaldehyde (Thermo Scientific; 28906), and after washing twice with PBS, stained with Hoechst33342 (Thermo Scientific; 62249) at 20uM. Nuclei were imaged by widefield microscopy and segmented using StarDist 2D with a ‘2D_versatile_fluo’ pretrained model. Segmented images were manually inspected to identify characteristics of debris and erroneously large segmentations, and size thresholding was used to remove these segmentations. Outlier total intensity of Hoechst signal within the nuclei were also removed. Nuclei counts were used as a proxy for cell counts, and by taking note of the total culture time and the final cell number, we calculated the cells’ doubling time.

### Size-dependent proteomics mass spectrometry

*P_TRE_ shCCND1* and *P_TRE_ CCND1* cells were incubated with doxycycline and harvested at days 0, 3, 4, 5, and 6 of induction. Samples were processed as below in ‘LC-MS/MS sample preparation’ and ‘TMT labeling for LC-MS/MS’. To sort cells by cell cycle, cells induced with doxycycline for 5 days were incubated with 20µM Hoechst33342 (Thermo Scientific; 62249) for 30 minutes at 37℃ before being sorted on a FACS Aria Fusion. Cells were sorted for G1 and G2 using the Hoechst stain for all 4 conditions and promptly processed for TMT mass spectrometry.

### LC-MS/MS sample preparation

Protein samples were prepared following a protocol adapted from a previous study.^35^ Briefly, cells were trypsinized and pelleted by centrifugation at 1000xG for 5 minutes and lysed for 30 minutes on ice in RIPA lysis buffer (Abcam; ab156034) with a protease and phosphatase inhibitor cocktail (Thermo Scientific; 78440). Lysates were cleared by centrifugation at 16000xG for 10 minutes at 4℃. Samples were then denatured by 1% SDS, reduced with 5mM DTT, alkylated with 10mM iodoacetamide, then precipitated with three volumes of 50% acetone and 50% ethanol. Proteins were solubilized with 2M urea, 50 mM Tris-HCl, pH 8.0, and 150mM NaCl, then digested with TPCK-treated trypsin (50:1) overnight at 37℃. Following digestion, peptides were acidified with trifluoroacetic acid and desalted with Sep-Pak 50mg C18 columns (Sep-Pak; WAT054955). The columns were pre-conditioned with 80% acetonitrile and 0.1% acetic acid, and washed with 0.1% trifluoroacetic acid. After loading the peptides, the columns were then washed with 0.1% acetic acid and eluted with 80% acetonitrile and 0.1% acetic acid. The eluents were dried in a concentrator at 45℃.

### Stable isotope labeling in cell culture and pulse chase

Prior to the beginning of the turnover experiment, *P_TRE_ shCCND1* and *P_TRE_ CCND1* cells were cultured in special lysine/arginine-free DMEM/F-12 media for SILAC (Thermo Scientific; 88370) supplemented with 10% dialyzed FBS (Gibco; 26400-044), 1% penicillin/streptomycin (Gibco; 15140-122), and ‘light’ versions of 0.499mM lysine (Cambridge Isotope Laboratories; ULM-8766-PK) and 0.699mM arginine (Cambridge Isotope Laboratories; ULM-8347-PK) for two weeks (approximately 14 doubling time). ‘Light’ version of proline (Sigma Aldrich; P0380-100G) was also added to a final concentration of 1.89mM to minimize the metabolic conversion of arginine to proline. Afterwards, the cells were induced with doxycycline to either knockdown or overexpress wildtype cyclin D1. To minimize the effects of cell crowding on growth rate, we cultured cells on a staggered schedule so that the cells were split every two days, and there were dishes ready to be collected every day of the collection phase. On the fifth day of induction, the media was switched to DMEM/F-12 with ‘heavy’ versions of lysine (Cambridge Isotope Laboratories; CNLM-291-H-1) and arginine (Cambridge Isotope Laboratories; CNLM-539-H-1) (same concentrations as light versions). Samples were collected for both cell lines after 0,1,3,6,12,24,50,120, and 240 hours and processed as described below in ‘LC-MS/MS sample preparation.’ For the biological replicate experiment, the cells were labeled and chased as above, but in reverse amino acid isotope order (from heavy to light).

### TMT labeling for LC-MS/MS

Following protein digestion and clean up, the peptides were labeled with TMT as adapted from Zecha et al. (2019) and the Thermo TMT10plex^TM^ Isobaric Label Reagent Set Protocol as described in Zatulovskiy et al. (2022).^35,56^ Briefly, 10-20µg of peptides were labelled with 100 µg of Thermo TMT10plex^TM^ for 1 hour. The reaction was quenched with 5% hydroxylamine for 15 minutes. Peptides were then pooled and acidified with 10% trifluoroacetic acid, desalted, and eluted with Sep-Pak C18 columns as described above in “LC-MS/MS sample preparation”.

### Peptide fractionation

Peptides were fractionated using a Pierce High pH Reversed-Phase Peptide Fractionation kit. Dried peptides were reconstituted in 0.1% TFA. Peptide concentrations were determined using a nanodrop before injection.

### LC/MS-MS data acquisition

TMT-labeled peptides were processed as described before.^33^ Briefly, they were resuspended in 0.1% formic acid and analyzed on a Fusion Lumos mass spectrometer (Thermo Fisher Scientific, San Jose, CA) equipped with a Thermo EASY-nLC 1200 LC system (Thermo Fisher Scientific, San Jose, CA). Peptides were separated by capillary reverse phase chromatography on a 25 cm column (75 µm inner diameter, packed with 1.6 µm C18 resin, AUR2-25075C18A, Ionopticks, Victoria Australia) and introduced with 180-min stepped linear gradient at a flow rate of 300 nL/min. The gradient steps were: 6–33% buffer B (0.1% (v:v) formic acid in 80% acetonitrile) for 145 min, 33-45% buffer B for 15 min, 40–95% buffer B for 5 min and maintain at 90% buffer B for 5 min. The column temperature was heated to 50 °C throughout the procedure. Xcalibur software was used for the data acquisition. Survey scans were acquired in the Orbitrap (centroid mode) over the range 380–1,400 m/z with a resolution of 120,000 (at m/z 200). For MS1, the normalized AGC target (%) was 250 and the maximum injection time was 100 ms. Ions were fragmented by collision-induced dissociation (CID) with normalized collision energies of 34. The isolation window was set to the 0.7-m/z window. MS2 was acquired in the ion trap mass analyzer with the scan rate set to ‘Rapid’, the normalized AGC target (%) was set to ‘Standard’, and maximum injection time to 35 ms. Dynamic exclusion of the sequenced peptides was set to 30 s. The maximum duty cycle time was set to 3 s. Relative changes in peptide concentration were determined at the MS3 level by isolating and fragmenting the five most dominant MS2 ion peaks.

### Spectral searches

All raw files were searched using the Andromeda engine embedded in MaxQuant (v2). Reporter ion MS3 search was conducted using TMT10-plex settings. Variable modifications were oxidation (M) and protein N-terminal acetylation. For pulse SILAC experiments, carbamidomethyl (C) was a fixed modification. The number of modifications per peptide was capped at five. Digestion was tryptic (proline-blocked). Database search used the UniProt Human proteome. The minimum peptide length was 7 amino acids. 1% FDR was determined using a reverse decoy proteome.

### Mass spectrometry data analysis

Peptides were first filtered out for decoy and contaminants. Each TMT channel was normalized by the total intensity in all TMT channels that were loaded together. Any peptides with the resulting ratio values 0 or less were filtered out. Each peptide intensity was normalized by the mean of all peptide intensities in each TMT channel. This value was used as a measure of relative concentration of the peptide in the given TMT channel. To get the change in relative concentration of the peptide with cell size, we transformed the concentration values of the peptide and the respective originating cell’s volumetric mean-normalized measurements by log2. We then drew a linear regression line using np.polyfit and obtained the slope value for that peptide. To find the change in concentration of a protein with cell size increase, the slope values for all peptides matching to that protein were averaged. Only proteins with at least two unique peptide matches were carried forward for further analysis. Peptides were differentiated by their modified sequence, charge, fraction of loading, and leading razor protein.

### Pulse SILAC-TMT data analysis

Peptides were filtered out for decoy, contaminants, and missed cleavage sites bearing both light and heavy versions of amino acids. Each peptide was uniquely identified by incorporating sequence, charge, fraction, oxidative state, N-terminal modifications, and leading razor protein information. Any peptides that still had duplicates were taken out of consideration. We adapted methods from Zecha et. al. (2018) to normalize the data.^56^ Firstly, any peptides that had missing entries at the highest theoretical maximum (first time point for amino acid being chased, last time point for amino acid being incorporated) were taken out, so did any peptides that had less than 3 timepoints from which to infer changing concentrations. Next, peptides bearing light versions of amino acids were matched with those bearing the heavy counterparts. If there were more than one matching pair, they were resolved by picking the peptide with the highest intensity. After this initial matching, additional peptide matches were made by considering a looser definition of unique peptide ID, dropping the fraction information. Reporter intensities of theoretically maximum TMT channels were normalized for both light and heavy amino acid-bearing peptides by n-row normalization as previously described.^56^ The following factor was derived and multiplied to all TMT channels for a given peptide (pep).

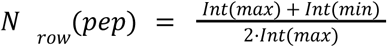

Where ‘Int(max)’ denotes the reporter intensity of peptide at the theoretically most abundant timepoint, and ‘Int(min)’ the opposite, the theoretically least abundant timepoint. In tracking chased light amino acids for example, ‘Int(max)’ would be timepoint 0, and ‘Int(min)’ would be the last timepoint. Next, a total sum normalization factor, N_sum_, was calculated:

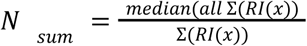

Where RI(X) is the reporter intensity channel X. This factor normalizes the mixing differences in TMT samples. Every TMT channel was multiplied by this factor.

### Protein half-life calculation

At steady-state, rates of protein incorporation equal the rates of turnover. Although we measured both rates, we chose to use only the turnover rate for simplicity. For every peptide, the intensity values were normalized by dividing by the theoretical maximum time point intensity value. For the isotope being chased, this was the first timepoint, and for the isotope being pulsed, the last timepoint. The resulting intensity values of all peptides belonging to a specific protein were fit to a kinetic model as described in Bosivert et. al. (2012) and Welle et al. (2016).^96,97^ Isotopes being chased were fit to the equation:

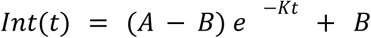

using optimize.curve_fit module from scipy, with input time variable in hours. K is the rate constant of turnover, A the maximum of the curve (ideally 1), and B the offset (ideally 0). Initial guesses for parameter searchers for K, A, B, were 0.05,1, and 0, with bounds set at [0,1],[0.5,1.5],[0,0.5] respectively. Half-life of the protein was found by solving the equation

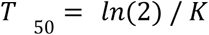

For quality control, the peptides were filtered such that R^2^ > 0.7, 0.67 < A < 1.3, -0.3 < B < 0.4. To get the T_50_ ratio between large and small cells for a given protein, the average T_50_ was taken for small and large cells across the replicates, and then divided as follows:

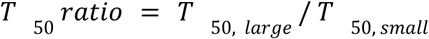

### Lysosome function assays

Cells were induced to different sizes as outlined above using the *P_TRE_ shCCND1* and *P_TRE_ CCND1* lines and doxycycline for 5 days. On day 5 of induction, the cells were induced with 3mM LLoMe (Santa Cruz Biotechnology; sc-285992A) or the equivalent volume of the carrier, ethanol. After 3 hours of treatment, the cells were trypsinized and spun down to remove the trypsin. After re-suspending the cells in growth media, the cells were allowed to recover at 37C for 30 min. Cells were then stained with LIVE/DEAD 405 marker (Thermo L34955; 0.1uL per 100k cells) and Annexin V 647 conjugate (Thermo A23204; 5uL per 100k cells) in Annexin binding buffer (Thermo; V13246) at room temperature for 30 min. Afterwards, the cells were washed once with Annexin binding buffer with 1% BSA (Fisher Scientific; BP1605-100), then re-suspended in the same buffer. The samples were immediately taken for analysis by flow cytometry. Laser power was adjusted to maximize separation of live and dead samples. Flow cytometry data was analyzed by FlowJo and all events were taken except for very small particulates indicating cell debris. The data was then gated to separate singlets and finally quantified to determine the percentage of dead and live cells, with the dead cells brightly fluorescent from LIVE/DEAD and Annexin V markers, and live cells negative for both. We note that the gates were similar across conditions, but manually adjusted for each sample due to the difference in cell size. Experiments involving chloroquine (Santa Cruz Biotechnology; sc-205629B) were analyzed in a similar manner, but focused on the most abundant single-cell cluster instead of using everything but cell debris.

For samples analyzed using microscopy, cells were similarly induced to new sizes, but applied a range of inhibitor doses starting on the 5th day of size-induction. For chloroquine, cells were treated for 0,1.56,3.125,6.25,12.5,25,50,100,200,400,800,1600 uM for 3 days. For bortezomib (Fisher Scientific; HY102275MG) treated cells, 0,0.625,1.25,2.5,5,10,20,40,80,160,320,640 nM were applied for 3 days. At the end of their treatment, cells were stained with 0.5uM SYTOX Deep Red Nucleic Acid (Invitrogen, S11380) and 10uM Hoechst33342 by directly adding concentrated versions of the reagents directly on top of the cells to minimize disturbance to the cells, then immediately analyzed by ImageXpress Micro Confocal system (Molecular Device) with widefield imaging using a 10x 0.45 Plan Apochromat 4mm WD objective. Cells were imaged with DAPI channel (excitation: 360/28 nm, emission: 447/60 nm, dichroic: 409 nm, exposure 50 ms, power: 10%) and Cy5 (excitation: 631/28 nm, emission: 692/40 nm, dichroic: 660 nm, exposure 100 ms, power: 10%). Images obtained were segmented for nuclei using the Hoechst33342 DNA stain with StarDist module with a pretrained ‘2D_versatile_fluo’ model, and SYTOX Deep Red stain quantified within the nuclei as a measure of cell death after subtracting background SYTOX signal. Signal concentration cutoff threshold to determine dead state was determined by a combination of gaussian mixture model separation and manual inspection. Proportion of dead cells for a given dose was determined and plotted against dosage.

### Cell cycle distribution analysis

Cell cycle distributions were measured by flow cytometry following staining of cells by 20µM Hoechst33342 for 30 minutes in 37C. Data was processed using FlowJo. After gating for singlets, the cells were separated by their DNA content and side scatter to distinguish G1 and G2M populations. Percentage of S phase populations were subsequently calculated by assuming that all cells were in one of G1, G2M, or S phases. For *HBEC-kt* cells, DNA was similarly stained, but imaged on a widefield epifluorescence microscope on a 10x objective. Nuclei were segmented using StarDist with a pretrained ‘2D_versatile_fluo’ model, and the background-subtracted total intensity of Hoechst signal used to measure DNA content. Proportion of G1 and G2M cells were distinguished as outlined above.

### RNA extraction

*P_TRE_ shCCND1* and *P_TRE_ CCND1* cells were induced with doxycycline for 4 or 5 days before sample collection. DMSO control samples were also prepared simultaneously. After PBS wash, cells were lysed using TRIReagent (Zymo Research; R2050-1-50), and RNA was extracted by Direct-zol^TM^ RNA Microprep Kit (Zymo Research; R2051-A).

### RNA-seq library preparation

mRNA was enriched using the NEBNext Poly(A) mRNA Magnetic Isolation Module (NEB, #E7490), and NEBNext Ultra II Directional RNA Library Prep Kit for Illumina (NEB, #E7760) was then used to prepare libraries for paired-end (2×150bp) Illumina sequencing (Novogene). More than 20 million reads were sequenced per sample.

### RNA-seq data processing

RNA-seq reads were aligned to the *H. sapiens* hg38 genome assembly and the transcriptome annotation from H. sapiens gene models from the v29 version of the GENCODE annotation (PMID: 30357393). For the purposes of RNA-seq data quality evaluation, genome browser track generation, reads were aligned against the genomes and set of splice junctions using the STAR aligner (version 2.5.3a; settings: --limitSjdbInsertNsj 10000000 --outFilterMultimapNmax 50 --outFilterMismatchNmax 999 --outFilterMismatchNoverReadLmax 0.04 --alignIntronMin 10 --alignIntronMax 1000000 --alignMatesGapMax 1000000 --alignSJoverhangMin 8 --alignSJDBoverhangMin 1 --sjdbScore 1 --twopassMode Basic --twopass1readsN -1) (PMID: 23104886). Read mapping statistics and genome browser tracks were generated using custom Python scripts. For quantification purposes, reads were aligned as 2×50mers in transcriptome space against an index generated from annotations described above using Bowtie (version 1.0.1; settings: -e 200 -a -X 1000) (PMID: 19261174). Alignments were then quantified using eXpress (version 1.5.1) (PMID: 23160280) before effective read count values and TPM (Transcripts Per Million transcripts) were quantified.

### mRNA slope value calculation

For transcriptomic data, the slope values were calculated similarly as it was done for the proteomics data, except using the TPM values themselves as proxy for relative concentrations. Only genes with more than 1 TPM were taken for further analysis. TPM values for a given gene from a particular day and replicate were normalized by the mean and log_2_ transformed. The mean of the median cell sizes of the samples taken the same day and replicate was used to normalize the cell sizes and similarly transformed by log_2_. Next, linear regression was performed using np.polynomial to obtain the slope value of the change in relative gene concentration with cell size. These slope values across all replicates and days were then averaged to find the mean slope value of the gene.

### Ordinary Least Squares Regression

statsmodels.api.OLS was used to build a predictive model for size-dependent proteome slopes. As the predictor variables, we used the mRNA slope and protein half-life measurements. For protein slope measures, proteins with more than 2 unique peptide mappings were used, and for protein half-life measurements, proteins with half-lives shorter than the estimated cell doubling time were used.

### SLAMseq cell culture

*P_TRE_ shCCND1* and *P_TRE_ CCND1* cells were induced with doxycycline for the entire duration of the experiment. On day 4 of induction, 10µM 4-thiouridine (s^4^U) (Selleck Chemicals; E1292) was added to the growth medium. A separate replicate was also made at the same time that instead received 10µM uridine (negative control)(Sigma Aldrich; U3003-5G). Media was exchanged every 3 hours for 24 hours. After 24 hours of s^4^U addition, cells were washed twice with PBS, and growth media with 1mM uridine was applied. At this time, the negative control and 0 hour timepoint samples were collected, and the chase was started. At each collection timepoint, the cells were washed once with PBS, and lysed with TRIReagent. Samples were collected at 0, 10, 20, 45, 90, 180, 360, 720, and 1440 minutes following initial media exchange with 1mM uridine. Cell size was measured for these samples on the same day.

### SLAMseq library preparation, sequencing, and data analysis

RNA quality control, library preparation, sequencing and initial data processing were performed by Lexogen NGS Services, Lexogen GmBH, Austria. In brief, samples were characterized by UV-Vis spectrophotometry (Nanodrop 2000c, ThermoFisher) and RNA integrity was assessed on a Fragment Analyzer System using the DNF-471 RNA Kit (15nt) (Agilent). The libraries were constructed using Lexogen’s QuantSeq 3’ mRNA-Seq Library Prep Kit FWD V2. RNA input to the library preparation was normalized to 75ng across all samples. Prepared libraries were quality controlled on a Fragment Analyzer System using the DNF-474 using the HS-DNA kit (1-6000bp) (Agilent). cDNA libraries were sequenced with an Illumina NovaSeq X platform using 100nt single end read length, with 40 million reads on average per sample. Data was analyzed using the SLAMdunk pipeline, as outlined on https://github.com/t-neumann/slamdunk/blob/bluebee-rc1/LICENSE

### mRNA data normalization for SLAMseq

Prior to further analysis, reads mapping to more than one gene were taken out. For all reads mapping to a given gene, thymine to cytosine conversion (T>C) rate was calculated by summing the number of T>Cs and dividing by the number of thymines in the reads. T>C conversion rate from the negative control was used as a measure of background noise and subtracted from all samples in a gene-specific manner. Any gene that had less than 1 read count per million was taken out. Genes with no T>C conversion in the first timepoint (theoretical maximum) were also taken out. Numbers of unique genes detected in samples collected between 0-180 min chase time were then examined, and any timepoint that had less than 60% of genes labelled compared to the number of genes detected with no T>C conversions (which formed a vast majority of reads detected) were omitted. Samples with no valid first timepoint were omitted outright. Next, only genes with at least 4 timepoints available to fit a decay curve were kept.

### mRNA half-life calculations

For each gene, T>C conversion rate of each timepoint was normalized by the first timepoint. Normalized T>C conversion rate for each gene were then fit on an exponential decay curve:

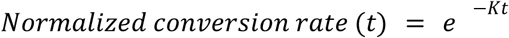

where K is the turnover rate, and t the timepoint of collection time. The half-life of the gene is

then:

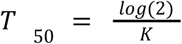

Genes with fits of R^2^ less than 0.6 were removed.

### High-throughput burst imaging

*Hbec-kt3* cells were seeded at densities ranging from 2 x 10^3^ to 1 x 10^4^ cells per well in a 96-well glass-bottom imaging plate 4 days prior to imaging. Fresh culture medium was applied the following day and replaced again on the day of imaging. Time-lapse imaging was performed overnight, capturing 8-12 fields of view per well. Imaging was conducted using high-throughput dual spinning disk microscopes (Yokogawa Cell Voyager 7000 or 8000). A 60x water immersion objective (1.2 NA) was used with a 488 nm excitation laser to quantify bursting activity and a 561 nm excitation laser for additional imaging (data from the 561 nm laser were not included in the final manuscript). A quad-band dichroic mirror (405/488/561/604 nm) was employed, and fluorescence was detected using a 16-bit Andor Neo 5.5 sCMOS camera with 2 x 2 pixel binning and a 525/50 nm bandpass emission filter. The XY pixel was 216.6 nm. For each field of view, a Z-stack of 16 images was acquired with a 0.5 um step size and maximum intensity projected for analysis. Time-lapse imaging was performed at 100-second intervals, with flat-field correction applied in real-time by the Yokogawa control software.

### Segmentation and tracking of burst

Each frame in the movie was normalized by csbdeep.utils.normalize with pmin=1. Next, nuclei were segmented using StarDist2D’s pretrained ‘2D_versatile_fluo’ model. Nuclear size distribution in the frame was inspected and aberrantly large and small nuclear segmentations were removed. Any nucleus on the frame borders was also removed. Nuclei were tracked using LapTrack with track_cost_cutoff = 50. Only tracks with more than 50 frames were kept. Bursts were segmented and tracked separately for each nucleus. For a given nucleus, any nuclear area change more than 25% between a single timepoint (100s) was considered erroneous segmentation and the entire nucleus was removed from analysis. For each frame, the nucleus was normalized from the raw image by csbdeep.utils.normalize with pmin=1, and bright spots identified by skimage.feature.blob_log with max_sigma=2. This module calculates the Laplacian of the Gaussian images with increasingly larger standard deviation. For each potential bright spot, a disk with radius=2 was drawn around it, and the intensity of the pixels summed. Background subtraction was done by subtracting the mean pixel value of the nucleus. Next, random positions inside the nuclei were selected and a disk with the same size as before was drawn. The region of the nucleus from which random positions were drawn was limited to avoid overlapping of disks drawn. Out of all possible positions, about a third were sampled. Intensity of pixels inside randomly selected and drawn disks were summed and similarly background subtracted. The standard deviation of the summed intensity inside these randomly selected disks was used to filter out low quality bright spots:

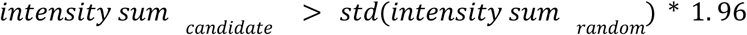

For ease of analysis, we set the shortest burst to be analyzed at 5 min. We also noted that bursts at some points significantly dimmed. We therefore considered for a given potential bright spot belonging to a burst, a 3-frame window in both time directions, and required that a bright spot have at least two neighboring bright spots. These bright spots also needed to be close enough spatially. After having selected higher quality bright spots, we then looked back at the original set of spots detected by the blob_log module and added back any spots that neighbor the high quality bright spots in time and space. From these spots, we then constructed tracks (bursting periods containing bright spots) from the tracking algorithm LapTrack. Only tracks longer than 2 frame lengths were kept and nuclei with less than 2 tracks throughout the entire movie were taken out entirely. If there were overlapping tracks, only the longest track was kept. For any timepoints before the first bright spot in a burst, random positions inside the nucleus were selected and logged as a proxy to the real location of the transcription site. A similar procedure was carried out for the timepoints between the last bright spot in the last burst and the last frame in the movie. For timepoints in between the bursting tracks, a linear interpolation was carried out to find a locus proxy for the real location of the tagged allele.

Background-subtracted intensity sums of all locations were then calculated. This now created one contiguous track for the entire length of the movie, consisting of bursting and inactive periods. The intensity sums across time were smoothened by a Savitzky-Golay filter with a window size of 5 and order 0. The individual bursting periods were then shortened based on the standard deviation of randomly detected spots in each frame (2.56x standard deviation of background). Remaining burst tracks were then allowed to expand using a looser threshold (1.96x standard deviation of background). Finally, all burst periods within one timeframe of another were merged.

### Bursting parameter acquisition

First and last instances of ON and OFF periods were discarded since we did not know their start and end times respectively. ON and OFF times were taken as the length of time during which we observed bursting or no bursting, respectively. We calculated burst size by taking the area under the curve of the background subtracted intensity sum of the bright spots and the time of bursting using sklearn.metrics.auc. Average nuclear area during the ON and OFF times were used as the cell size during these times.

## Code availability

Codes used to process all data are available at https://github.com/cdsyou/You_et_al_2025

## Acknowledgments

We thank Georgi Marinov for technical assistance in analyzing RNAseq data. We thank Shuyuan Zhang for assistance and feedback on various aspects of experiments covered in this paper. Evgeny Zatulovskiy gave numerous, detailed feedback to this paper, for which we are grateful. We also thank Shicong Xie for assistance with implementing Python-related data analysis. We thank the Stanford Shared FACS facility for assistance in cell sorting using instruments purchased by the Parker Institute for Cancer Immunotherapy (NIH S10 Shared Instrument Grant). Finally, we thank the Sarafan ChEM-H High-Throughput Screening Knowledge Center for assistance in imaging lysosome inhibition assays (SIG S10OD026899).

## References

1. Aarts, P. A. M. M., Bolhuis, P. A., Sakariassen, K. S., Heethaar, R. M. & Sixma, J. J. Red Blood Cell Size Is Important for Adherence of Blood Platelets to Artery Subendothelium. Blood 62, 214–217 (1983).

2. Snyder, G. K. & Sears, R. D. Red blood cell size and the Fåhraeus–Lindqvist effect. Can. J. Zool. 84, 419–424 (2006).

3. Sharp, D. S. et al. Mean Red Cell Volume as a Correlate of Blood Pressure. Circulation 93, 1677–1684 (1996).

4. Farnier, C. et al. Adipocyte functions are modulated by cell size change: potential involvement of an integrin/ERK signalling pathway. Int. J. Obes. 27, 1178–1186 (2003).

5. Hatton, I. A., et al. The human cell count and size distribution. Proc. Natl. Acad. Sci. 120, e2303077120 (2023).

6. Lloyd, A. C. The Regulation of Cell Size. Cell 154, 1194–1205 (2013).

7. Ginzberg, M. B., Kafri, R. & Kirschner, M. On being the right (cell) size. Science 348, 1245075–1245075 (2015).

8. Jorgensen, P. & Tyers, M. How Cells Coordinate Growth and Division. Curr. Biol. 14, R1014–R1027 (2004).

9. Sandlin, C. W. et al. Epithelial cell size dysregulation in human lung adenocarcinoma. PLOS ONE 17, e0274091 (2022).

10. Nguyen, A., Yoshida, M., Goodarzi, H. & Tavazoie, S. F. Highly variable cancer subpopulations that exhibit enhanced transcriptome variability and metastatic fitness. Nat. Commun. 7, 11246 (2016).

11. Li, Q., Rycaj, K., Chen, X. & Tang, D. G. Cancer stem cells and cell size: A causal link? Semin. Cancer Biol. 35, 191–199 (2015).

12. Davies, D. M., van den Handel, K., Bharadwaj, S. & Lengefeld, J. Cellular enlargement - A new hallmark of aging? Front. Cell Dev. Biol. 10, (2022).

13. Lengefeld, J. et al. Cell size is a determinant of stem cell potential during aging. Sci. Adv. 7, eabk0271 (2021).

14. Demidenko, Z. N. & Blagosklonny, M. V. Growth stimulation leads to cellular senescence when the cell cycle is blocked. Cell Cycle 7, 3355–3361 (2008).

15. Ogrodnik, M. Cellular aging beyond cellular senescence: Markers of senescence prior to cell cycle arrest in vitro and in vivo. Aging Cell 20, e13338 (2021).

16. Schmoller, K. M. The phenomenology of cell size control. Curr. Opin. Cell Biol. 49, 53–58 (2017).

17. Zatulovskiy, E. & Skotheim, J. M. On the Molecular Mechanisms Regulating Animal Cell Size Homeostasis. Trends Genet. 36, 360–372 (2020).

18. Zatulovskiy, E., Zhang, S., Berenson, D. F., Topacio, B. R. & Skotheim, J. M. Cell growth dilutes the cell cycle inhibitor Rb to trigger cell division. Science 369, 466–471 (2020).

19. Zhang, S., Zatulovskiy, E., Arand, J., Sage, J. & Skotheim, J. M. The cell cycle inhibitor RB is diluted in G1 and contributes to controlling cell size in the mouse liver. Front. Cell Dev. Biol. 10, (2022).

20. Schmoller, K. M., Turner, J. J., Kõivomägi, M. & Skotheim, J. M. Dilution of the cell cycle inhibitor Whi5 controls budding-yeast cell size. Nature 526, 268–272 (2015).

21. Schmoller, K. M. et al. Whi5 is diluted and protein synthesis does not dramatically increase in pre-Start G1. Mol. Biol. Cell 33, lt1 (2022).

22. D’Ario, M. et al. Cell size controlled in plants using DNA content as an internal scale. Science 372, 1176–1181 (2021).

23. Liu, D., Lopez-Paz, C., Li, Y., Zhuang, X. & Umen, J. Subscaling of a cytosolic RNA binding protein governs cell size homeostasis in the multiple fission alga Chlamydomonas. PLOS Genet. 20, e1010503 (2024).

24. Keifenheim, D. et al. Size-Dependent Expression of the Mitotic Activator Cdc25 Suggests a Mechanism of Size Control in Fission Yeast. Curr. Biol. 27, 1491–1497.e4 (2017).

25. Chen, Y., Zhao, G., Zahumensky, J., Honey, S. & Futcher, B. Differential Scaling of Gene Expression with Cell Size May Explain Size Control in Budding Yeast. Mol. Cell 78, 359–370.e6 (2020).

26. Williamson, D. H. & Scopes, A. W. The distribution of nucleic acids and protein between different sized yeast cells. Exp. Cell Res. 24, 151–153 (1961).

27. Crissman, H. A. & Steinkamp, J. A. RAPID, SIMULTANEOUS MEASUREMENT OF DNA, PROTEIN, AND CELL VOLUME IN SINGLE CELLS FROM LARGE MAMMALIAN CELL POPULATIONS. J. Cell Biol. 59, 766–771 (1973).

28. Liu, X., Oh, S. & Kirschner, M. W. The uniformity and stability of cellular mass density in mammalian cell culture. Front. Cell Dev. Biol. 10, (2022).

29. Berenson, D. F., Zatulovskiy, E., Xie, S. & Skotheim, J. M. Constitutive expression of a fluorescent protein reports the size of live human cells. Mol. Biol. Cell 30, 2985–2995 (2019).

30. Lanz, M. C. et al. Increasing cell size remodels the proteome and promotes senescence. Mol. Cell 82, 3255–3269.e8 (2022).

31. Mäkelä, J. et al. Genome concentration limits cell growth and modulates proteome composition in Escherichia coli. eLife 13, (2024).

32. Cheng, L. et al. Size-scaling promotes senescence-like changes in proteome and organelle content. 2021.08.05.455193 Preprint at 10.1101/2021.08.05.455193 (2021).

33. Lanz, M. C. et al. Genome dilution by cell growth drives starvation-like proteome remodeling in mammalian and yeast cells. Nat. Struct. Mol. Biol. 31, 1859–1871 (2024).

34. Tan, C., Lanz, M. C., Swaffer, M., Skotheim, J. & Chang, F. Intracellular diffusion in the cytoplasm increases with cell size in fission yeast. Mol. Biol. Cell 36, ar51 (2025).

35. Zatulovskiy, E. et al. Delineation of proteome changes driven by cell size and growth rate. Front. Cell Dev. Biol. 10, (2022).

36. Miettinen, T. P. & Björklund, M. Cellular Allometry of Mitochondrial Functionality Establishes the Optimal Cell Size. Dev. Cell 39, 370–382 (2016).

37. Neurohr, G. E. et al. Excessive Cell Growth Causes Cytoplasm Dilution And Contributes to Senescence. Cell 176, 1083–1097.e18 (2019).

38. Khurana, A., Chadha, Y. & Schmoller, K. M. Too big not to fail: Different paths lead to senescence of enlarged cells. Mol. Cell 83, 3946–3947 (2023).

39. Crozier, L. et al. CDK4/6 inhibitor-mediated cell overgrowth triggers osmotic and replication stress to promote senescence. Mol. Cell 83, 4062–4077.e5 (2023).

40. Foy, R. et al. Oncogenic signals prime cancer cells for toxic cell overgrowth during a G1 cell cycle arrest. Mol. Cell 83, 4047–4061.e6 (2023).

41. Manohar, S. et al. Genome homeostasis defects drive enlarged cells into senescence. Mol. Cell 83, 4032–4046.e6 (2023).

42. Wilson, G. A. et al. Active growth signaling promotes senescence and cancer cell sensitivity to CDK7 inhibition. Mol. Cell 83, 4078–4092.e6 (2023).

43. Mu, L. et al. Mass measurements during lymphocytic leukemia cell polyploidization decouple cell cycle- and cell size-dependent growth. Proc. Natl. Acad. Sci. 117, 15659–15665 (2020).

44. Ginzberg, M. B. et al. Cell size sensing in animal cells coordinates anabolic growth rates and cell cycle progression to maintain cell size uniformity. eLife 7, e26957 (2018).

45. Devany, J., Falk, M. J., Holt, L. J., Murugan, A. & Gardel, M. L. Epithelial tissue confinement inhibits cell growth and leads to volume-reducing divisions. Dev. Cell 58, 1462–1476.e8 (2023).

46. Saftig, P. & Klumperman, J. Lysosome biogenesis and lysosomal membrane proteins: trafficking meets function. Nat. Rev. Mol. Cell Biol. 10, 623–635 (2009).

47. Yang, C. & Wang, X. Lysosome biogenesis: Regulation and functions. J. Cell Biol. 220, e202102001 (2021).

48. Bajaj, L. et al. Lysosome biogenesis in health and disease. J. Neurochem. 148, 573–589 (2019).

49. Curnock, R. et al. TFEB-dependent lysosome biogenesis is required for senescence. EMBO J. 42, e111241 (2023).

50. López-Otín, C., Blasco, M. A., Partridge, L., Serrano, M. & Kroemer, G. The Hallmarks of Aging. Cell 153, 1194–1217 (2013).

51. Hernandez-Segura, A., Nehme, J. & Demaria, M. Hallmarks of Cellular Senescence. Trends Cell Biol. 28, 436–453 (2018).

52. Mathieson, T. et al. Systematic analysis of protein turnover in primary cells. Nat. Commun. 9, 689 (2018).

53. Schwanhäusser, B. et al. Global quantification of mammalian gene expression control. Nature 473, 337–342 (2011).

54. Doherty, M. K., Hammond, D. E., Clague, M. J., Gaskell, S. J. & Beynon, R. J. Turnover of the Human Proteome: Determination of Protein Intracellular Stability by Dynamic SILAC. J. Proteome Res. 8, 104–112 (2009).

55. Sabatier, P. et al. Global analysis of protein turnover dynamics in single cells. Cell 188, 2433–2450.e21 (2025).

56. Zecha, J. et al. Peptide Level Turnover Measurements Enable the Study of Proteoform Dynamics. Mol. Cell. Proteomics 17, 974–992 (2018).

57. Buccitelli, C. & Selbach, M. mRNAs, proteins and the emerging principles of gene expression control. Nat. Rev. Genet. 21, 630–644 (2020).

58. Liu, Y., Beyer, A. & Aebersold, R. On the Dependency of Cellular Protein Levels on mRNA Abundance. Cell 165, 535–550 (2016).

59. Vogel, C. & Marcotte, E. M. Insights into the regulation of protein abundance from proteomic and transcriptomic analyses. Nat. Rev. Genet. 13, 227–232 (2012).

60. Swaffer, M. P. et al. RNA polymerase II dynamics and mRNA stability feedback scale mRNA amounts with cell size. Cell 186, 5254–5268.e26 (2023).

61. Herzog, V. A. et al. Thiol-linked alkylation of RNA to assess expression dynamics. Nat. Methods 14, 1198–1204 (2017).

62. Tani, H. et al. Genome-wide determination of RNA stability reveals hundreds of short-lived noncoding transcripts in mammals. Genome Res. 22, 947–956 (2012).

63. Raj, A., Peskin, C. S., Tranchina, D., Vargas, D. Y. & Tyagi, S. Stochastic mRNA Synthesis in Mammalian Cells. PLOS Biol. 4, e309 (2006).

64. Rodriguez, J. & Larson, D. R. Transcription in Living Cells: Molecular Mechanisms of Bursting. Annu. Rev. Biochem. 89, 189–212 (2020).

65. Miller, O. L. & McKnight, S. L. Post-replicative nonribosomal transcription units in D. melanogaster embryos. Cell 17, 551–563 (1979).

66. Miller, O. L. & Beatty, B. R. Visualization of Nucleolar Genes. Science 164, 955–957 (1969).

67. Berry, S., Müller, M., Rai, A. & Pelkmans, L. Feedback from nuclear RNA on transcription promotes robust RNA concentration homeostasis in human cells. Cell Syst. S2405471222001703 (2022) doi:10.1016/j.cels.2022.04.005.

68. Zhurinsky, J. et al. A Coordinated Global Control over Cellular Transcription. Curr. Biol. 20, 2010–2015 (2010).

69. Sun, X.-M. et al. Size-Dependent Increase in RNA Polymerase II Initiation Rates Mediates Gene Expression Scaling with Cell Size. Curr. Biol. 30, 1217–1230.e7 (2020).

70. Padovan-Merhar, O. et al. Single Mammalian Cells Compensate for Differences in Cellular Volume and DNA Copy Number through Independent Global Transcriptional Mechanisms. Mol. Cell 58, 339–352 (2015).

71. Golding, I., Paulsson, J., Zawilski, S. M. & Cox, E. C. Real-Time Kinetics of Gene Activity in Individual Bacteria. Cell 123, 1025–1036 (2005).

72. Chubb, J. R., Trcek, T., Shenoy, S. M. & Singer, R. H. Transcriptional Pulsing of a Developmental Gene. Curr. Biol. 16, 1018–1025 (2006).

73. Larson, D. R., Zenklusen, D., Wu, B., Chao, J. A. & Singer, R. H. Real-Time Observation of Transcription Initiation and Elongation on an Endogenous Yeast Gene. Science 332, 475–478 (2011).

74. Suter, D. M. et al. Mammalian Genes Are Transcribed with Widely Different Bursting Kinetics. Science 332, 472–474 (2011).

75. Bothma, J. P. et al. Dynamic regulation of eve stripe 2 expression reveals transcriptional bursts in living Drosophila embryos. Proc. Natl. Acad. Sci. 111, 10598–10603 (2014).

76. Wan, Y. et al. Dynamic imaging of nascent RNA reveals general principles of transcription dynamics and stochastic splice site selection. Cell 184, 2878–2895.e20 (2021).

77. Berenson, D. F., Zatulovskiy, E., Xie, S. & Skotheim, J. M. Constitutive expression of a fluorescent protein reports the size of live human cells. Mol. Biol. Cell 30, 2985–2995 (2019).

78. Tunnacliffe, E. & Chubb, J. R. What Is a Transcriptional Burst? Trends Genet. 36, 288–297 (2020).

79. Patange, S. et al. MYC amplifies gene expression through global changes in transcription factor dynamics. Cell Rep. 38, 110292 (2022).

80. Trzaskoma, P. et al. 3D chromatin architecture, BRD4, and Mediator have distinct roles in regulating genome-wide transcriptional bursting and gene network. Sci. Adv. 10, eadl4893 (2024).

81. Pomp, W., Meeussen, J. V. W. & Lenstra, T. L. Transcription factor exchange enables prolonged transcriptional bursts. Mol. Cell 84, 1036–1048.e9 (2024).

82. Gressel, S., Schwalb, B. & Cramer, P. The pause-initiation limit restricts transcription activation in human cells. Nat. Commun. 10, 3603 (2019).

83. Liu, X., Kraus, W. L. & Bai, X. Ready, pause, go: regulation of RNA polymerase II pausing and release by cellular signaling pathways. Trends Biochem. Sci. 40, 516–525 (2015).

84. Core, L. & Adelman, K. Promoter-proximal pausing of RNA polymerase II: a nexus of gene regulation. Genes Dev. 33, 960–982 (2019).

85. Vidal, P. J., Pérez, A. P., Yahya, G. & Aldea, M. Transcriptomic balance and optimal growth are determined by cell size. Mol. Cell 84, 3288–3301.e3 (2024).

86. Lessenger, A. T., Skotheim, J. M., Swaffer, M. P. & Feldman, J. L. Somatic polyploidy supports biosynthesis and tissue function by increasing transcriptional output. J. Cell Biol. 224, e202403154 (2024).

87. López-Otín, C., Blasco, M. A., Partridge, L., Serrano, M. & Kroemer, G. Hallmarks of aging: An expanding universe. Cell 186, 243–278 (2023).

88. Liu, S. et al. Oversized cells activate global proteasome-mediated protein degradation to maintain cell size homeostasis. eLife 14, e75393 (2025).

89. Xie, S., Swaffer, M. & Skotheim, J. M. Eukaryotic Cell Size Control and Its Relation to Biosynthesis and Senescence. Annu. Rev. Cell Dev. Biol. 38, annurev-cellbio-120219-040142 (2022).

90. Davies, D. M., van den Handel, K., Bharadwaj, S. & Lengefeld, J. Cellular enlargement - A new hallmark of aging? Front. Cell Dev. Biol. 10, (2022).

91. Chadha, Y., Khurana, A. & Schmoller, K. M. Eukaryotic cell size regulation and its implications for cellular function and dysfunction. Physiol. Rev. 104, 1679–1717 (2024).

92. Fassl, A. et al. Increased lysosomal biomass is responsible for the resistance of triple-negative breast cancers to CDK4/6 inhibition. Sci. Adv. 6, eabb2210 (2020).

93. Huang, X. et al. Prolonged early G1 arrest by selective CDK4/CDK6 inhibition sensitizes myeloma cells to cytotoxic killing through cell cycle–coupled loss of IRF4. Blood 120, 1095–1106 (2012).

94. Chan, K. Y., et al. GPX4-dependent ferroptosis sensitivity is a fitness trade-off for cell enlargement. iScience 28, (2025).

95. Meerbrey, K. L. et al. The pINDUCER lentiviral toolkit for inducible RNA interference in vitro and in vivo. Proc. Natl. Acad. Sci. 108, 3665–3670 (2011).

96. Welle, K. A. et al. Time-resolved Analysis of Proteome Dynamics by Tandem Mass Tags and Stable Isotope Labeling in Cell Culture (TMT-SILAC) Hyperplexing*. Mol. Cell. Proteomics 15, 3551–3563 (2016).

97. Boisvert, F.-M. et al. A Quantitative Spatial Proteomics Analysis of Proteome Turnover in Human Cells*. Mol. Cell. Proteomics 11, M111.011429 (2012).

